# Integrated ‘omics analysis reveals human milk oligosaccharide biosynthesis programs in human lactocytes

**DOI:** 10.1101/2025.03.17.643803

**Authors:** Sarah Kate Nyquist, Laasya Devi Annepureddy, Kristija Sejane, Annalee Furst, G Devon Trahan, Michael C Rudolph, Alecia Jane Twigger, Lars Bode, Barbara E Engelhardt, Jayne F Martin Carli, Britt Anne Goods

## Abstract

Human milk oligosaccharides (HMOs) are integral to infant health. Yet, their complex biosynthesis pathways in the mammary gland during lactation remain under characterized. To address this knowledge gap, we performed integrated analyses of single-cell RNA-sequencing (scRNA-seq) datasets combined with select HMO concentration measures. We identify differential expression patterns of known HMO synthesis genes in epithelial subsets and nominate several candidate genes that vary with HMO concentration. Additionally, we identify novel gene patterns and transcription factors that may regulate the expression of HMO biosynthesis genes and the cellular pathways supporting HMO production. Finally, we demonstrate that co-expression of HMO synthesis genes and milk fat synthesis genes is limited, suggesting distinct epithelial cell subtypes may be responsible for the production of different milk components. Our study suggests that HMO synthesis may be achieved through cell type specialization within the lactocyte compartment.

## Introduction

Human milk oligosaccharides (HMOs) are unconjugated glycans produced in the mammary gland during lactation. HMOs act as prebiotics and antimicrobials in the infant’s gut, and they may also directly modulate infant intestinal epithelial and immune cell responses.^1–6^ HMO concentrations in human milk are highest in colostrum (20-25 g/L) and subsequently decline during mature lactation (5-20 g/L). On average, total HMO concentration is higher than total milk protein concentrations (8 g/L), making HMOs a substantial reservoir of bioactive stimuli present during postnatal development.^7^ These biomolecules exhibit incredible structural diversity, with over 150 different types in human milk.^8,9^

Maternal genotype appears to play a primary role in HMO composition with evidence for variants in a single gene, fucosyltransferase 2 (*FUT2*), splitting lactating people into two groups— secretors and non-secretors— based on the presence or absence of some fucosylated HMOs. The profile of these and other individual HMOs is highly consistent following successive pregnancies in individual women.^10^ Additional maternal and infant-related factors, such as maternal diet, feeding practices, and environmental factors, are also thought to determine HMO profiles, including maternal diet, feeding practices, and environmental factors.^11,12^ Production of HMOs occurs through sequential enzymatic reactions involving glycosyltransferases, which transfer specific sugar residues to the growing oligosaccharide chain (**Figure 1A**). It is generally thought that all secretory mammary epithelial cells in the lactating mammary gland (lactocytes) synthesize HMOs during lactation, however, this has not been directly investigated and HMO-producing lactocytes are poorly classified. Insights into HMO biosynthesis may inform the development of interventions to improve infant health outcomes, particularly in preterm infants and in settings where breastfeeding is not feasible or is challenging.

**Figure 1:**
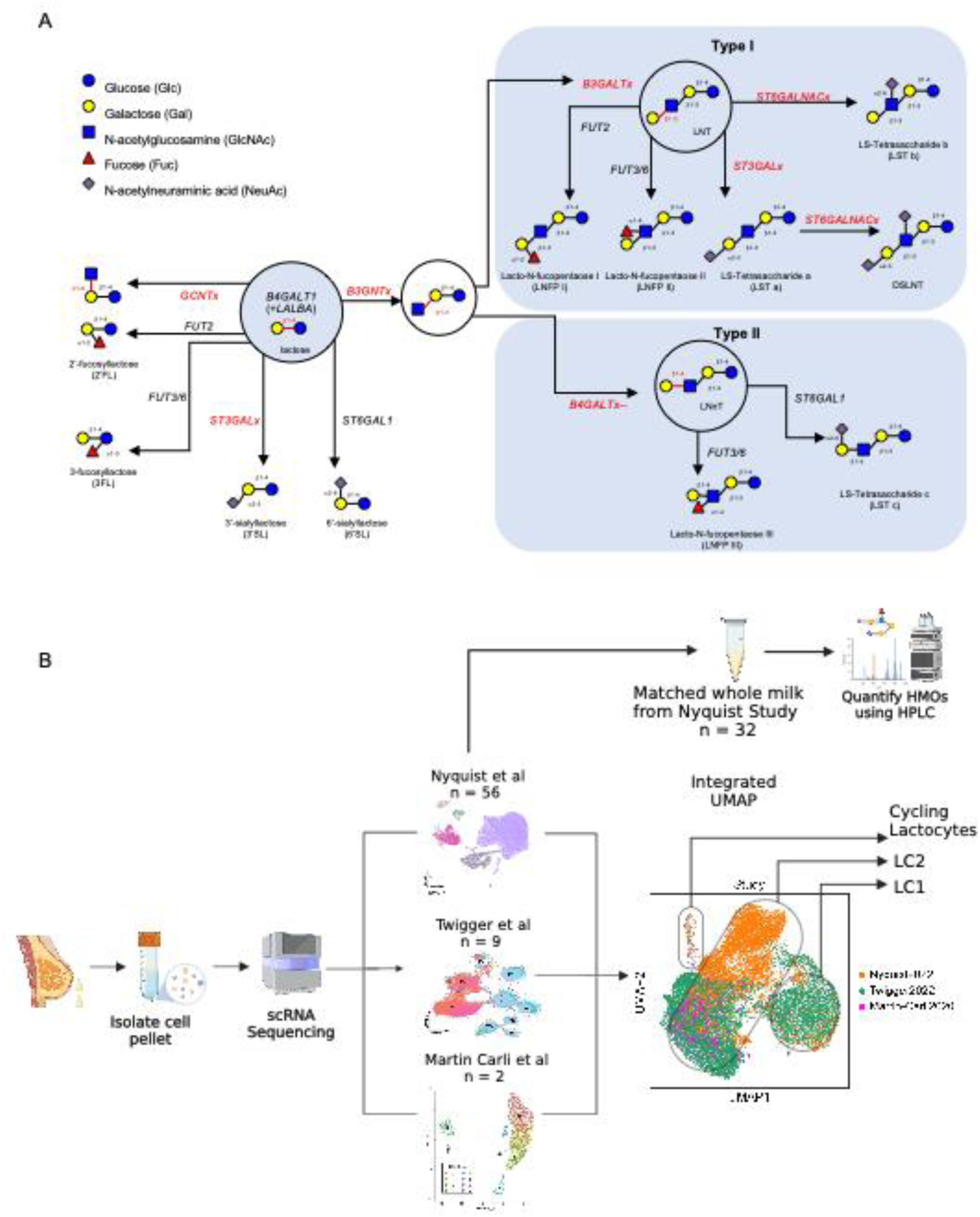
Overview of HMO hypothesized biosynthesis pathways and study design. A. HMO biosynthesis network previously described in the literature, including candidate genes or gene families responsible for synthesis steps, beginning with lactose which is synthesized by glycosyltransferase *B4GALT1* in conjunction with lactalbumin alpha (*LALBA*). Genes with x suffixes indicate unknown/uncertain enzymes in the family. Type I and type II groupings are based on angle of beta 1 addition of galactose at the lactose reducing end. B. Workflow of the study using existing scRNA datasets, depicted as UMAPs, with n = number of samples. Larger integrated UMAP produced using Harmony. Glycomic profiles generated from matched whole milk samples to Nyquist study scRNA-seq samples using high performance liquid chromatography (HPLC). Created with BioRender.com.

Despite their importance to infant health, little is understood about HMO biosynthesis in humans and the regulation of HMO diversity in human milk. This largely due to limitations in experimental methods and models for studying lactating human cells. Robust cell culture models of the human lactating mammary gland do not exist resulting in a barrier to understanding of how lactocytes create essential components and transport them into human milk. *In-vivo* models also have limitations as the milk oligosaccharides found in murine or other mammalian milk differ from the human-specific HMOs.^13^ Given these challenges, the full complement of HMO biosynthetic genes has not yet been defined. Genomic data-driven approaches, however, hold the potential to tease apart possible routes of HMO biosynthesis in human lactocytes by leveraging correlative analyses in milk samples to identify genes involved in the biosynthesis of HMOs. Extensive computational modeling of bulk data identified *FUT2*, ST6 beta-galactoside alpha-2,6-sialyltransferase 1 (*ST6GAL1*), fucosyltransferase 3 (*FUT3*), and beta-1,its 4-galactosyltransferase 4 (*B4GALT4*) genes involved in HMO biosynthesis, showing promise for further transcriptomic studies.^14^ Still, bulk RNA-seq analysis limits the characterization of the cell types involved in these processes, which is essential given the known heterogeneity in lactocytes. A higher resolution understanding of genes, cofactors, and cellular processes correlated with HMO synthesis in subsets of lactocytes would improve our understanding of HMO production and the complex regulation of HMOs in the human mammary gland.

To this end, several single-cell RNA-seq (scRNA-seq) datasets of cells shed into human milk have characterized the cell types in the lactating mammary.^15–19^ These datasets have consistently identified two major epithelial populations, called lactocyte type 1 (LC1) and lactocyte type 2 (LC2) previously referred to as luminal clusters 1 and 2. LC1 cells express genes involved in cellular organization and epithelial barrier maintenance, key for establishing tight junctions and polarity in epithelial cells, in addition to milk production genes. LC2 cells are characterized as a secretory cell type that express genes involved in milk production, secretion, and synthesis of macromolecules, peptides, and lipids. Milk-derived LC1 and LC2 cells are generally considered representative of lactocytes *in vivo*, although this has yet to be formally established. To date, no studies have leveraged the scRNA-seq data from these cells to deeply explore how these two cell types may contribute to HMO biosynthesis.

In this study, we combine scRNA-seq data and HMO concentrations quantified from matched samples to identify HMO synthesis pathway genes associated with distinct LC1 and LC2 cell types. We leverage three independent scRNA-seq studies of human milk-derived cells to map HMO synthesis gene expression onto the cell types found in human milk. When combined, the overall number of cells and total number of samples create a powerful resource to study the complex process of HMO production, overcoming the lack of power within each discrete study. Next, we integrate RNA and HMO concentration data to identify glycosyltransferases, transporters, and other cellular processes correlated with HMO concentrations to gain insight into the cell types, processes, and genes that regulate HMO biosynthesis. Taken together, our work suggests differences in mammary HMO biosynthetic capacity between LC1 and LC2 cells during lactation, nominates putative genes and pathways that support HMO production, and overall suggests the potential for specification of lactocytes in producing different milk constituents.

## Results

### Previously identified HMO synthesis genes have different expression profiles in LC1 and LC2 cells

We first sought to better delineate epithelial cell subsets producing HMOs in the mammary gland during lactation and to determine whether their gene expression patterns are distinct among MEC subtypes. To accomplish this, we integrated three publicly available scRNA-seq human milk cell datasets and identified the subsets of epithelial cells, including LC1 and LC2 cells, expressing genes implicated in HMO synthesis, including the well-characterized lactalbumin alpha (*LALBA*) and beta-1,4-galactosyltransferase 1 (*B4GALT1*).^20^ By harmonizing these datasets, we identified reproducible gene expression across multiple collection sites and milk-derived cell types. In total, our integrated dataset includes 111,257 single cells from 71 samples (**Figure 1B**). Shared cell types were identified across the merged datasets, including immune cells and LC1 and LC2 lactocyte cell clusters, which are thought to be responsible for the production of HMOs.

With this integrated resource, we first confirmed the expression of validated and hypothesized HMO synthesis genes, such as enzyme encoding genes *ST6GAL1* and *FUT2.*^13^ We also investigated several candidate gene families whose functions in other oligosaccharide-producing tissues suggest they may be involved in HMO synthesis, including fucosyltransferases (e.g., fucosyltransferase 6, *FUT6*), glucosaminyl (N-acetyl) transferases (e.g., glucosaminyl (N-acetyl) transferase 1, *GCNT1*), galactosyltransferases (e.g., beta-1,3-galactosyltransferase 4, *B3GALT4)*, acetylglucosaminyltransferases (e.g., UDP-GlcNAc:betaGal beta-1,3-N-acetylglucosaminyltransferase 4, *B3GNT4*), and sialyltransferases (e.g., ST3 beta-galactoside alpha-2,3-sialyltransferase 1, *ST3GAL1*) (**Figure 1A, Supplementary Table 1**). We filtered this list by selecting genes that have RNA expression in lactocytes isolated from human milk (**Supplementary Table 1)** and found that most candidate HMO synthesis genes were more highly expressed in LC1 cells across samples and datasets. However, some were highly expressed in LC2 cells including *ST6GAL1* and *B3GALT4* (**Figure 2A**). *B4GALT1,* the gene encoding the catalytic subunit of lactose synthase, was highly expressed in both LC1 and LC2 cells.

**Figure 2:**
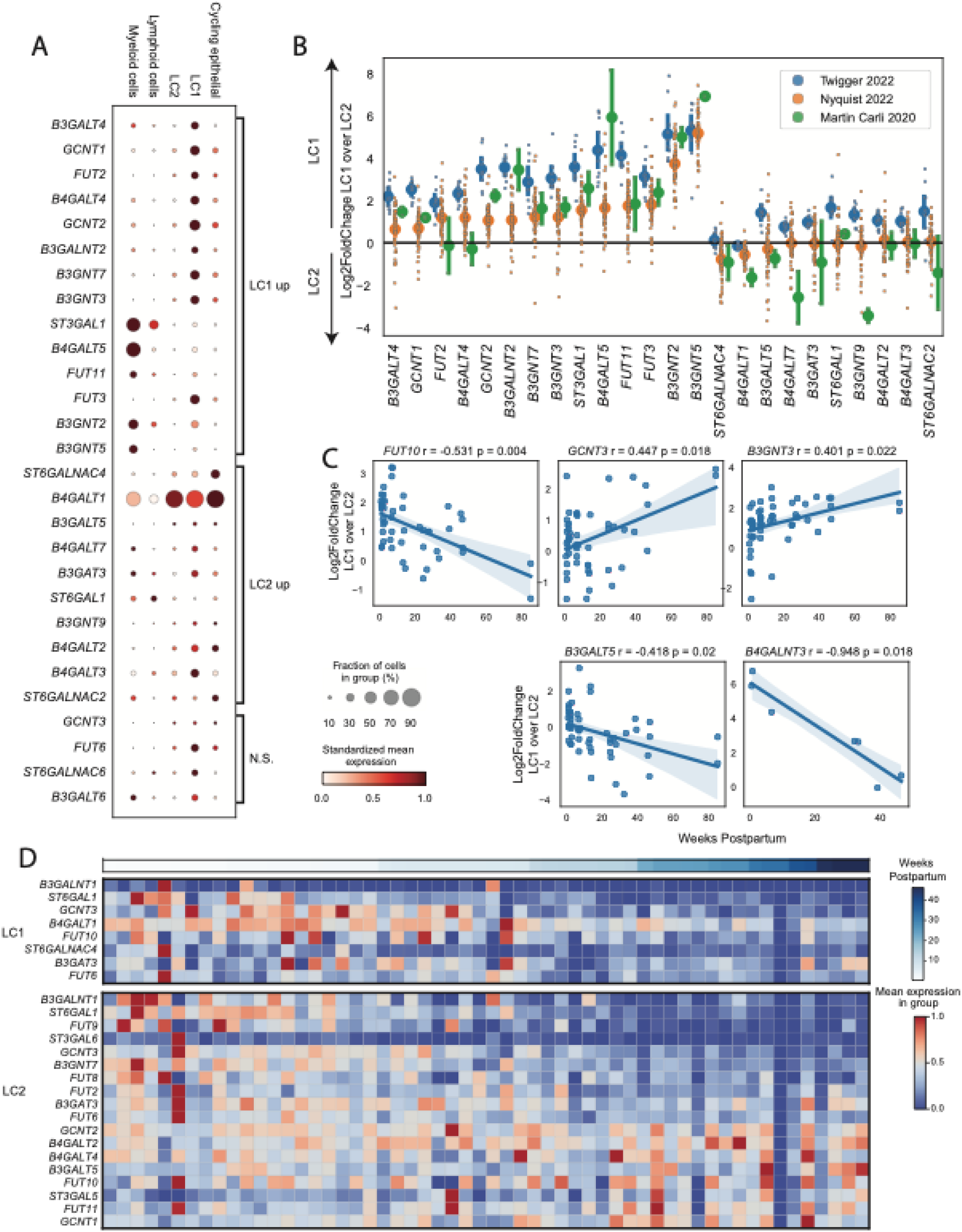
HMO synthesis genes are expressed in lactocytes isolated from human milk. A. Dotplot showing expression of select candidate HMO synthesis genes expressed in a minimum of 10 percent of epithelial cells (left) in integrated scRNA-seq data across studies grouped by major cell types (top). LC1 and LC2 enrichment of each gene identified using DESeq2 comparisons of pseudo bulk data between LC1 and LC2 cells. N.S. genes have large effect size but do not reach significance in the comparison. Dot size indicates the percent of cells in the group expressing each gene, and dot color indicates the column-standardized mean expression of each gene across cell type groups. B. Log2-fold change of pseudo bulk gene expression of genes significantly differentially expressed between LC1 and LC2 cells across studies. Each biological sample is represented by a small dot, larger dots indicate mean log2-fold change within each study, and colors indicate individual studies. Error bars indicate the 95% confidence interval around the mean. C. Spearman correlation of the log2-FoldChange between expression in LC1 cells and LC2 cells of potential HMO synthesis genes with time postpartum. P-values from BH corrected spearman correlations. Lines represent linear regression with confidence interval. D. Candidate HMO synthesis genes identified in Nyquist et al 2022 as associated with time postpartum. Heatmap includes z-scored mean expression of pseudo bulk data within each sample’s LC1 (top) or LC2 (bottom) population ordered by time postpartum.

Several candidate HMO genes are also expressed in non-lactating mammary gland tissue. Using data that also profiled breast tissue, we compared gene expression of these genes in resting mammary gland tissue to cells from milk. We found that, of the genes highly expressed in LC1 or LC2 cells, glucosaminyl (N-acetyl) transferase 2 (I blood group) (*GCNT2*), beta-1,3-N-acetylgalactosaminyltransferase 2 (*B3GALNT2*), UDP-GlcNAc:betaGal beta-1,3-N-acetylglucosaminyltransferase 2 (*B3GNT2*), beta-1,3-glucuronyltransferase 3 (*B3GAT3*), beta-1,4-galactosyltransferase 3 (*B4GALT3*), and beta-1,5-galactosyltransferase 6 (*B4GALT6*) were also highly expressed in luminal progenitor (LP) or hormone-responsive (HR) cells in the non-lactating mammary gland (**Supplementary Figure 1A**).^16^

To further describe lactocyte subtype differences in the expression of HMO synthesis genes between LC1 and LC2 cells and the consistency of these differences across samples and studies, we calculated the log2-fold change of normalized gene expression between LC1 and LC2 cells within each sample. We found candidate genes whose expression was higher in LC1s than in LC2s, and this difference was reproducible across samples and studies. For genes more highly expressed in LC2s, the log2-fold change between LC1 and LC2 expression had less overall agreement across studies and samples (**Figure 2B**).

Next, because these samples were collected at different times postpartum, and because HMO concentration is known to vary across this timeframe, we determined whether any of these expression patterns varied with the duration of lactation for the LC1 or LC2 cells. Our prior work identified many changes in gene expression in both LC1 and LC2 cells over time postpartum, including several potential HMO biosynthesis genes.^17^ In particular, *FUT2* decreases in expression over time postpartum in LC2 cells (**Figure 2C**). Overall, the expression of most candidate HMO production genes decreases with time postpartum in both LC1 cells and LC2 cells. This is consistent with the previously described decrease in overall HMO concentration in human milk with time postpartum.^21^

Given that the expression of these genes varies over time postpartum and the log2-fold changes of the differences in these genes’ expression between LC1 and LC2 cells were variable across samples and studies, we asked if their difference in expression between LC1 and LC2 cells was also associated with time postpartum. To do this, we quantified the amount of variability in our integrated dataset that is explained by time postpartum. We found that fucosyltransferase 10 (*FUT10*), beta-1,3-galactosyltransferase 5 (*B3GALT5*), and beta-1,4-N-acetyl-galactosaminyltransferase 3 (*B4GALNT3*) were more highly expressed in LC1s at earlier time points and more highly expressed in LC2s at later time points (Pearson correlation Benjamini-Hochberg [BH] adjusted p-value ≤ 0.05). Conversely, glucosaminyl (N-acetyl) transferase 3, mucin type (*GCNT3*), and UDP-GlcNAc:betaGal beta-1,3-N-acetylglucosaminyltransferase 3 (*B3GNT3*) were more highly expressed in LC2s at earlier time points and more highly expressed in LC1s at later time points (Pearson correlation BH adjusted p-value ≤ 0.05), with many other genes trending toward associations with time not reaching significance (**Supplementary Figure 1B**, **Figure 2D)**. These may represent gene expression programs that are coordinately regulated as the mammary gland transitions from early to mature lactation

### Co-expression patterns of HMO synthesis genes in LC1 and LC2 cells reveal putative pathways of specific HMO production

To identify potential pathways responsible for the synthesis of specific HMOs, we performed a co-expression analysis of genes expressed in LC1 or LC2 cells. Similar patterns of co-expression of certain genes can be used to infer their roles in the same cellular processes.^22,23^ Using a hypergeometric test on binarized expression values, we identified pairs of potential HMO production genes in LC1 and LC2 cells that are co-expressed within the same single cells more frequently than random given their total extent of expression across each study and sample (hypergeometric test p≤0.05). To quantify the reproducibility of this co-expression, we identified the percentage of samples for which there was adequate expression to power the comparison in which each pair of genes was co-expressed (**Figure 3A and B**). We found that the pairs of co-expressed genes in LC1 cells differed from those in LC2 cells and that more gene pairs showed co-expression in LC2 cells than in LC1 cells. A higher percentage of samples showed co-expression for specific pairs of genes in LC1 cells. Our analysis identified several candidate genes for the sialyl-lacto-N-tetraose b (LSTb) synthesis pathway that are co-expressed in both LC1 and LC2 cells, including genes UDP-GlcNAc:betaGal beta-1,3-N-acetylglucosaminyltransferase 7 (*B3GNT7*) or *B3GNT3*, *B3GALT5* or *B3GALT4*, ST6 N-acetylgalactosaminide alpha-2,6-sialyltransferase 2 (*ST6GALNAC2)* or 6 (*ST6GALNAC6)* (**Figure 3B**). We also identified candidate genes for the lacto-N-fucopentaose III (LNFPIII) synthesis pathway that are co-expressed in LC2 cells, including *B3GNT3* or *B3GNT7*, *B4GALT3* or beta-1,4-galactosyltransferase 2 (*B4GALT2*), and *FUT6* or fucosyltransferase 3 (*FUT3*).

**Figure 3:**
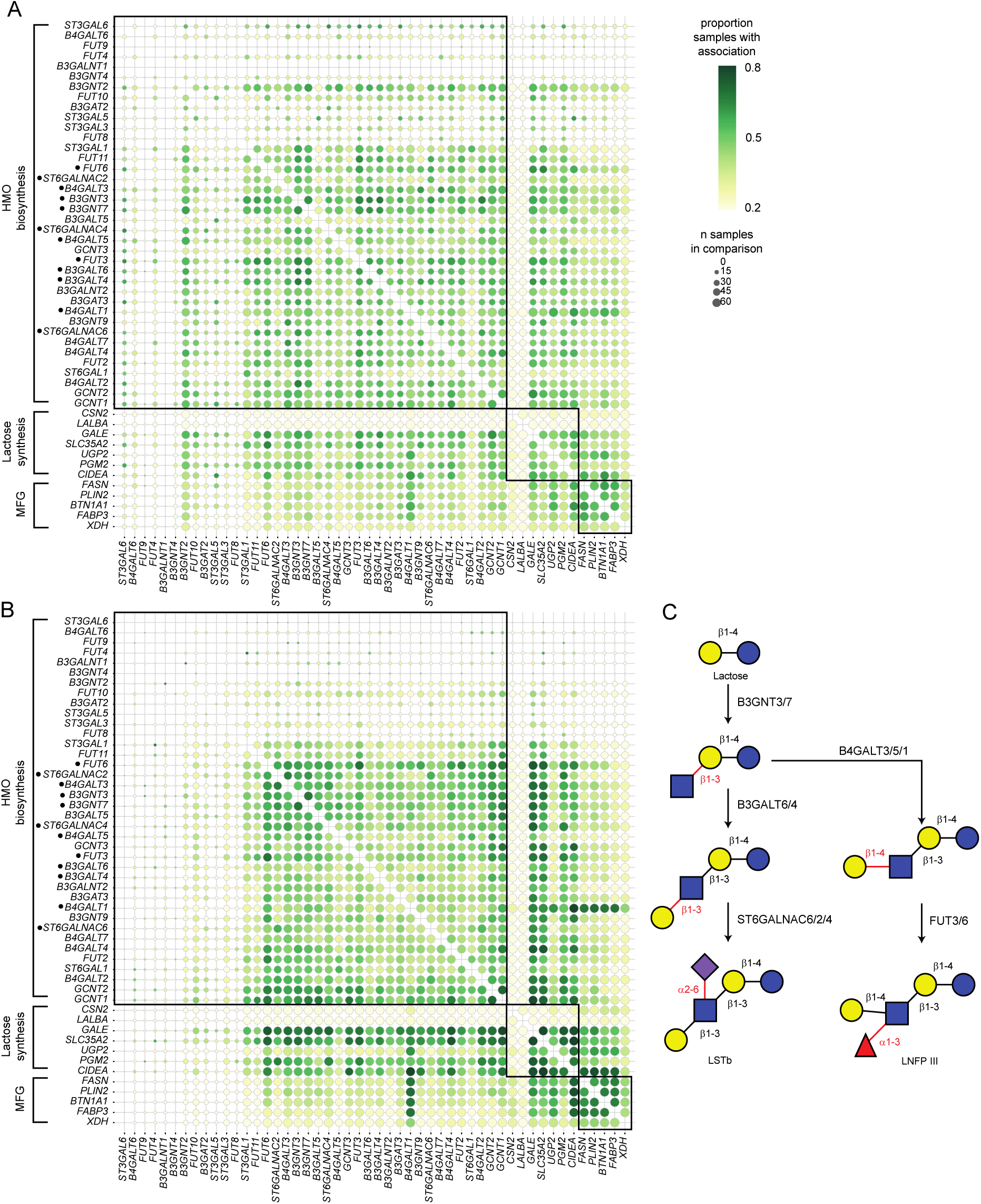
Co-expression analysis of candidate HMO synthesis genes nominates potential pathways. A. Candidate HMO synthesis genes, known casein and lactose synthesis genes, and known milk fat globule production genes co-expressed via hypergeometric test of pairwise co-occurrence of gene expression greater than 0 in single cells in LC1 (A) and LC2 (B) cell types across samples in all three integrated studies. Dot size indicates the number of samples for which sufficient cells express both genes to allow robust comparison, and color indicates percent of those eligible samples for which gene co-expression was more frequent than random. Genes are hierarchically clustered using WPGMC algorithm as implemented in scipy on the percent of samples associated (color in these plots). C. Candidate pathway for LSTb synthesis for which component genes *B3GNT7 or 3, B3GALT6 or 4, ST6GALNAC2 or 6* are identified as co-expressed in both LC1 and LC2 cell types across samples and for the LNFPIII pathway with component genes B4GALT3, 4, or 1 and FUT 3 or 6 (starred genes).

To contextualize and validate this gene-gene co-expression analysis, we considered the co-expression of genes involved in pathways known to be important for the synthesis of other human milk components including milk fat globule (MFG) production, casein and lactose synthesis, as well as glycolysis (**Figure 3A-B**). As expected, the majority of genes in these pathways show high levels of shared co-expression across all 71 samples in our integrated dataset. Two exceptions are the genes casein beta (CSN2) and *LALBA*, which are widely expressed across all lactocytes but do not appear to be co-expressed with many other genes in this set, demonstrating an edge case to this analysis: expression in all cells implies that a gene is not co-expressed because there is no variation in expression to correlate with other genes (**Figure 3A-B**).

In addition to using this approach to identify genes that are consistently co-expressed across samples, we can identify groups of genes that are consistently not co-expressed across samples. Our analysis revealed an interesting grouping in co-expressed genes. In particular, genes known to be involved in milk fat globule production (e.g., fatty acid synthase (*FASN*), fatty acid binding protein 3 *(FABP3),* and perilipin 2 *(PLIN2*)) are consistently co-expressed in both LC1 and LC2 cells. *B4GALT1*, which is common to both LC1 and LC2 cells, is strongly co-expressed with known lipid metabolism genes (*FASN*), cell death inducing DFFA like effector a (*CIDEA*)*, FABP3*, butyrophilin subfamily 1 member A1 (*BTN1A1*)*, PLIN2*, xanthine dehydrogenase (*XDH*), while other HMO synthesis genes were more frequently co-expressed with each other and not with milk metabolism genes among LC1 and LC2 cells (**Figure 3**). Similarly, genes known to be involved with HMO biosynthesis are consistently co-expressed in both LC1s and LC2s. However, genes across HMO and milk fat biosynthesis processes are not frequently co-expressed (**Figure 3A, B**). This suggests that the cell types that perform HMO biosynthesis may be distinct from those responsible for milk fat globule production, or these processes may be differentially regulated within these cell types.

### HMO concentrations vary temporally and correlate with cell type proportions

We next determined if paired HMO concentration data and scRNA-seq could be used to infer specific cell types that are important for HMO production according to their cell type proportions. To this end, we quantified HMO concentration using banked whole milk samples that matched the previously generated scRNA-seq samples. This analysis included 27 samples from 10 donors collected between 4 and 272 days postpartum (**Figure 4A, S2A, Table S2**). All milk donors included were secretors based on expression of the *FUT2* gene. Using a linear mixed model (LMM), we found that the concentration of most HMOs decreased with time, including 2′-fucosyllactose (2’FL), 3’-sialyllactose (3’SL), 6’-sialyllactose (6’SL), lacto-N-tetraose (LNT), lacto-N-neotetraose (LNnT), lacto-N-fucopentaose I (LNFPI), LNFPIII, LSTb, sialyl-lacto-N-tetraose c (LSTc), difucosyl-lacto-N-tetraose (DFLNT), lactose-N-hexaose (LNH), disialyl-lacto-N-tetraose (DSLNT), fucosyl-lacto-N-hexaose (FLNH), difucosyl-lacto-N-hexaose (DFLNH), fucosyl-disialyl-lacto-N-hexaose (FDSLNH) (BH-adjusted p-value ≤ 0.05; **Table S3; Figure 4B, S2B**). One HMO, 3-fucosyllactose (3FL), increased over time postpartum. These findings are consistent with other longitudinal studies of HMO concentrations over time postpartum.^21^

**Figure 4:**
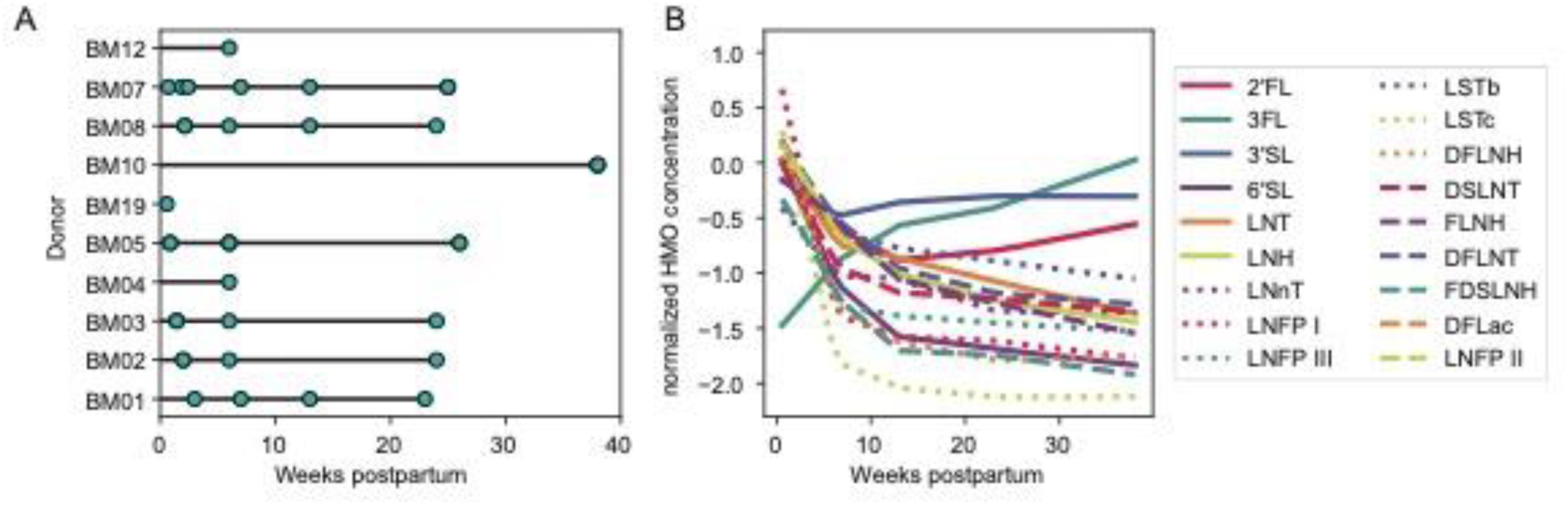
HMO concentrations have temporal variation and correlate with epithelial and immune cell type proportions. A. Samples with HMO concentration data across different times postpartum. Teal samples have matched scRNA-seq data from Nyquist et al 2022. B. Loess fits for association between each HMO concentration and time postpartum in samples from (A). HMOs with significant (BH adjusted p <= 0.05) association via linear mixed models included in this plot.

Next, using an LMM to account for time postpartum, we tested the association between LC1 and LC2 cell proportions in each milk sample with matched HMO concentration data. The only HMO associated with lactocyte cell proportions was LNT, which was positively associated with LC2 cell abundance (LMM BH-adjusted p-value ≤ 0.01; **Figure S2C, Table S3**). HMOs are also known to have immunomodulatory effects and to interact with immune cells such as macrophages and dendritic cells.^3,24^ Thus, we tested if any of the different HMO concentrations vary with immune cell populations present in breast milk. Dendritic cell abundance was negatively associated with 3’SL, LNT, LNFPIII, LSTb, LNH, DSLNT, and FLNH (BH-adjusted p ≤0.05; **Figure S2D**). HMOs 2’FL and DSLNT were positively associated with B cell proportions (BH-adjusted p ≤ 0.01), and HMOs, LNnT, LNFPI, lacto-N-fucopentaose II (LNFPII), and LSTb, were negatively associated with B cell proportions (BH-adjusted p ≤ 0.01). LNT, LSTb, and DFLNT concentrations were all negatively associated with neutrophil proportions (BH-adjusted p ≤ 0.05). Overall, we found little association between lactocyte type abundance and HMO concentration, but several associations within the immune cell compartment. This may indicate a shared interaction between the immune system and HMO production with a specific pathogen or other foreign exposure.

### Gene patterns associated with HMO concentrations are distinct between LC1 and LC2 subsets in matched samples

While there was little association between cell type abundance of LC1 and LC2 epithelial cell subsets and sample-level HMO concentrations, we hypothesized that the gene expression programs in LC1s and LC2s relevant to HMO production may be associated with matched HMO concentration. To test this, we used DESeq2 on pseudobulk gene expression counts of each sample’s LC1 and LC2 cells to identify genes potentially associated with sample-level HMO concentrations.^25^ We found hundreds of genes associated with the concentration of several HMOs (**Figure 5A, Table S4**). Gene expression in LC2s was associated with 2’FL, 3’SL, 6’SL, DFLNH, DSLNT, disialyl-lacto-N-hexaose (DSLNH), DFLNT, LNFPIII, and LSTc, while gene expression in LC1 cells was associated with LNFPI, FDSLNH, LSTb, and LNH. Few genes were shared between HMOs like LNFPI, DSLNT, and LNFPII, suggesting unique gene programs in these cell types for each HMO tested. (**Figure 5A, S3**) These results suggest that, while lactocyte abundance may not be associated with HMO concentration, there are genes within these LC1 and LC2 lactocytes that are associated with HMO concentration.

**Figure 5:**
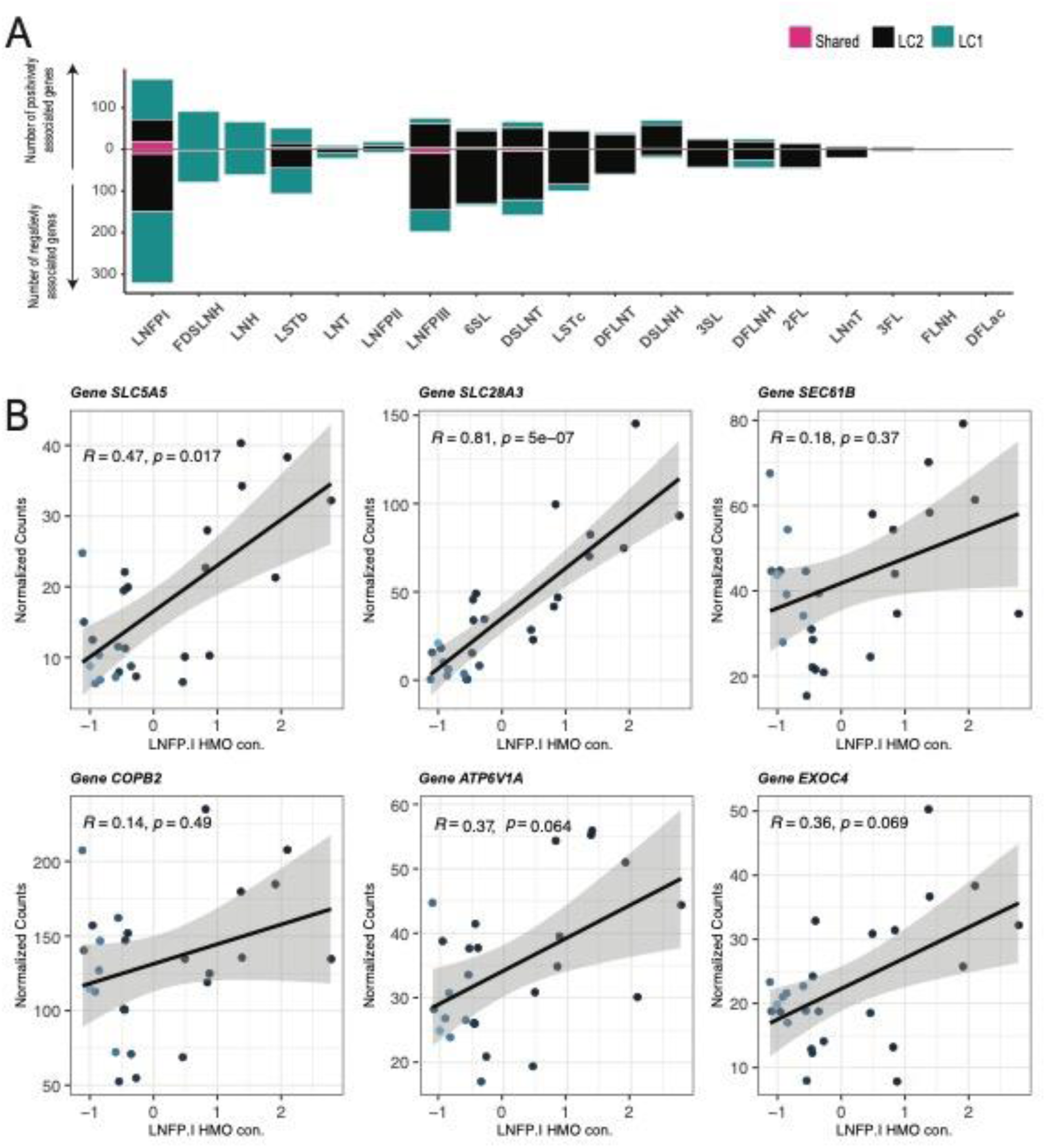
Gene patterns associated with HMO concentrations are distinct between LC1 and LC2 subsets in matched samples. A. Stacked bar plot of the number of positively associated (positive counts) and negatively associated (negative counts) genes (DESeq2, BH adjusted p <= 0.05) that are present in LC1 only (teal) and LC2 (black) only cell types as well as shared between LC1 & LC2 (pink). B. Linear regression between normalized counts of select transporter and enzyme genes (DESeq2 BH adjusted p-values < 0.05) associated with normalized LNFPI concentration in LC1 cells with 95% confidence intervals. (See also table S5)

We recovered associations of known glycosyltransferases involved in HMO biosynthesis. Notably, LC1s showed a positive association of *ST6GAL1* with 6’SL (BH-adjusted p-value ≤ 0.08), which has been previously characterized in *St6gal1*^−/−^ knockout mice that showed the absence of 6’SL oligosaccharide in their milk (**Table 1, Table S4)**.^13^ Instead of *ST6GAL1*, in LC2s we found a positive association of *ST3GAL1* expression with increasing concentrations of 6’SL, which was not expected given prior results. For other HMOs, LSTb, DSLNT, DFLNT, LNFPI, and LNFPIII—we identified positive associations with the expression of *ST6GAL1* in LC1 cells; LC2 cells, in contrast, exhibited positive associations with these HMOs and expression of *ST3GAL1* (**Table 1**). We observed a negative association of *B3GNT3* with the same HMOs (LSTb, DSLNT, DFLNT, LNFPI, and LNFPIII) in LC1s, and a negative association of *B3GALT4* in LC2s with DSLNT, LNFPI, and LNFPIII (**Table 1**). We also observed distinct gene expression patterns between LC1 and LC2 cells across various HMOs, indicating potentially different sets of genes associated with the production of these glycans within these cell types. Overall, the association of potential HMO synthesis genes, both positive and negative, with specific HMOs may suggest complex regulatory feedback, based on enzyme and substrate availability, between LC1 and LC2 cells and HMOs.

**Table 1:**
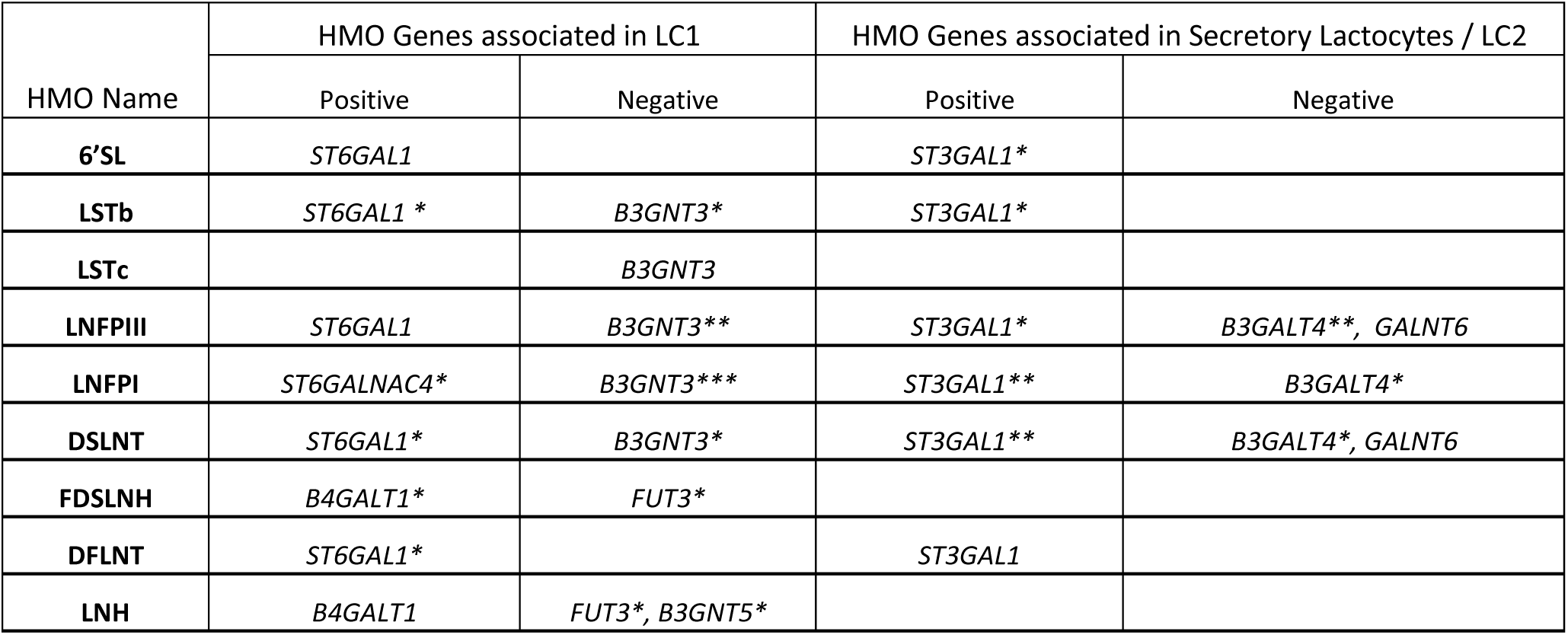
Positive and Negative associations of known HMO synthesis genes in LC1 and LC2 cells. The stars indicate different levels of statistical significance calculated from p-adjusted (BH) values using DESEQ-2 analysis. (See also Table S5) *padj <= 0.05, **padj <= 0.01, ***padj <= 0.001, no star indicates padj <= 0.1.

In addition to examining glycosyltransferase genes, our analysis identified pathways that may support HMO synthesis directly or indirectly. We annotated differentially expressed genes by their molecular characteristics to identify enzymes and transporters involved in HMO synthesis and transport through the ER-Golgi network.^26,27^ In LC1 cells, we found a positive association between sodium-dependent transporters known to be highly expressed in the lactating mammary gland and LNFPI concentration, specifically the sodium-iodide symporter solute carrier family 5 member 5 (*SLC5A5*) and the sodium-dependent nucleoside transporter solute carrier family 28 member 3 (*SLC28A3*) (**Figure 5B**).^28,29^ Additionally, a positive association was also noted between the protein transport protein SEC61 translocon subunit beta (*SEC61B*) and Golgi membrane proteins like COPI coat complex subunit beta 2 (*COPB2*) and ATPase H+ transporting V1 subunit A (*ATP6V1A*) with increasing LNFPI concentration (**Figure 5B**). The SEC61 translocon directs proteins to the ER, assisting in their binding and folding.^30^ Proteins are then packaged in vesicles and transported to the Golgi, where retrograde transport and sorting of glycosylation enzymes largely depend on COPI-coated vesicles.^31,32^ We also observed that the exocyst complex component 4 (*EXOC4*), involved in the vesicle transport machinery of secretory pathways, was associated with the concentration of LNFPI (**Figure 5B**).^33^ Our findings support the involvement of the ER-Golgi network in the transport of proteins involved in HMO biosynthesis. The genes outlined above have not previously been associated with HMO transport, and further validation is required.

### Analysis of gene patterns associated with Type-I and Type-II HMO concentrations reveal differences in transport and lipid synthesis processes in LC1 and LC2 cells

HMO synthesis is a structured process where some steps are expected to be shared between HMOs. HMOs are categorized into two groups based on their glycosidic linkages: Type-I HMOs, characterized by *β*1-3 linkage of galactose to N-acetyl-glucosamine, and Type-II HMOs, distinguished by *β*1-4 linkage of galactose (**Figure 1B**). We next tested the hypothesis that there may be shared processes used by LC1 and LC2 cells based on the type of HMO being produced. To do this, we used a similar approach as above, testing for gene expression correlated with the concentration of aggregate Type-I or Type-II HMOs. Like individual HMOs, our aggregate HMO analysis revealed an increased number of genes associated with Type-I or Type-II HMO concentration in the LC2 cells compared to the LC1 cells (**Figure S4, Supplementary Table 4**).

In LC2 cells, we saw a positive correlation of *ST3GAL1,* with increasing concentration of Type-I HMOs (BH adjusted p-value < 0.05) (**Figure 6A**). *ST3GAL1* is known to catalyze the transfer of sialic acid onto lactose or Gal*β*1-3-GalNAc-terminated glycoconjugates through a *α*2-3 linkage.^34,35^ This could potentially be involved in the synthesis of LSTa, the substrate to produce DSLNT. Also, with the concentration of Type-I HMOs, we identified a positive correlation with STT3B oligosaccharyltransferase complex (*STT3B*) and UDP-N-Acetylglucosamine (UDP-GlcNAc) pyrophosphorylase 1, (*UAP1*), involved in the synthesis of UDP-GlcNAc. (BH-adjusted p-value ≤ 0.05) (**Figure 6A**). UDP-GlcNAc is a precursor molecule for the biosynthesis of N-acetylglucosamine (GlcNAc)-containing glycans, which are abundant in HMOs. These are essential for the transfer of GlcNAc moieties onto the growing oligosaccharide chains during HMO biosynthesis.^36^ In the LC1 cells, we identified a negative association of (*B3GNT3)* between the concentration of Type-I HMOs. (**Figure 6A**) A family of *B3GNTx* enzymes is involved in the addition of GlcNAc at *β*2-3 linkage to galactose, as required at the initial stage in the production of Type-I and Type-II HMOs (**Figure 1A**). Therefore, further experiments are needed to verify the association of these enzyme transcripts and understand their protein products’ relationship to HMO synthesis.

**Figure 6:**
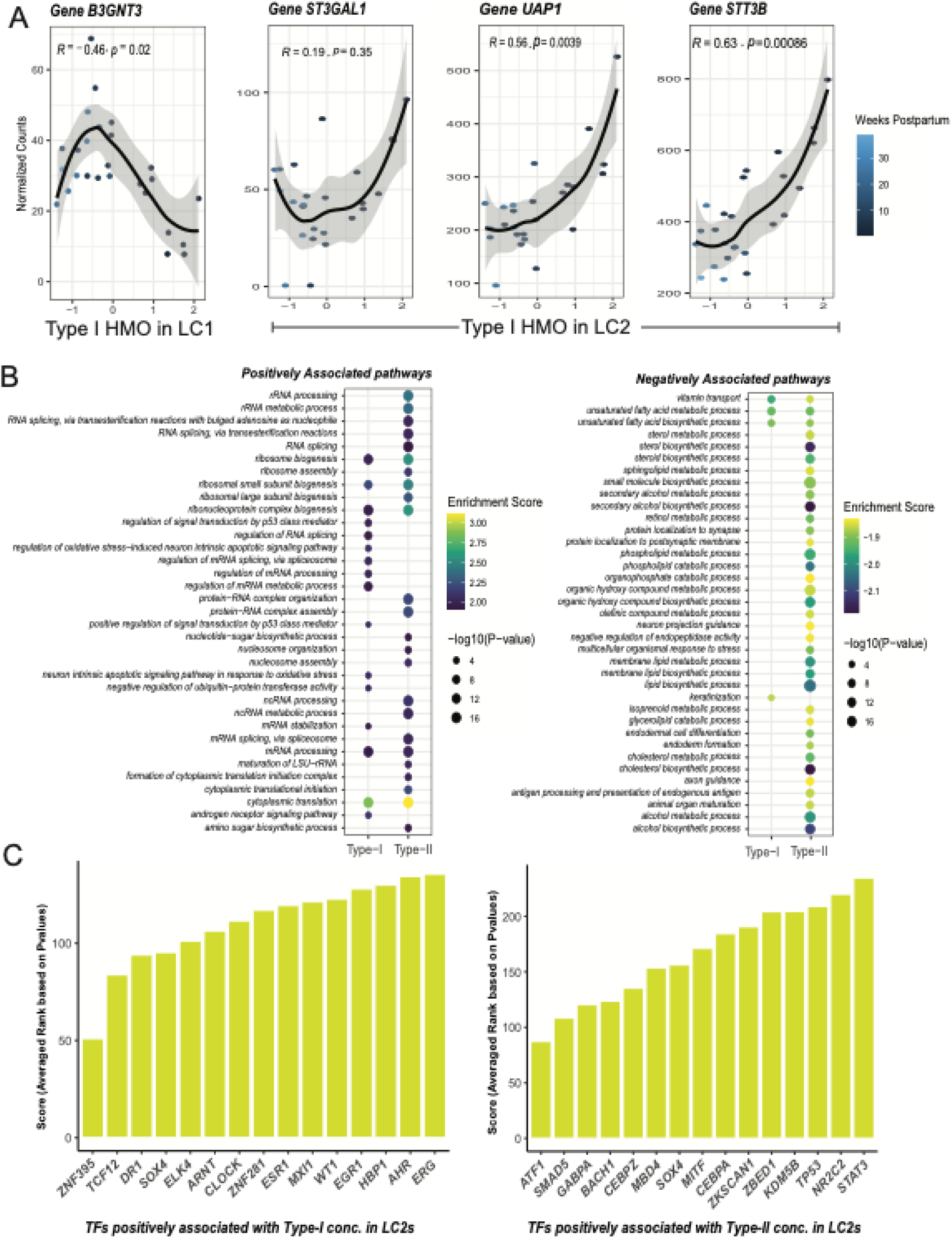
Type I HMO concentrations are associated with biosynthesis genes and pathways. A. Scatter Plot of the normalized Type-I HMO concentration versus normalized counts of select glycosyl transferases from DeSeq2 analysis in LC1s and LC2s. Adjusted p-value (BH) <0.05 **B.** GSEA GO BP dotplot of top 40 positively enriched pathways (right) and top 40 negatively enriched pathways (left) associated with Type I or Type II HMO concentrations in LC2. FDR < 0.05 **C.** Top 15 transcription factors associated with upregulated DEGs associated with Type-I HMO concentration in LC2 from CHEA analysis. (right) Top 15 positive transcription factors associated with upregulated DEGs associated with Type-II HMO concentration in LC2 from CHEA analysis. (right) The mean scores are averaged ranks of each TF across multiple gene sets in the CHEA database. The rank of each TF in a gene set is assigned based on BH adjusted p-values. Lower p-values (higher significance) indicate lower integer ranks and lower overall scores (see methods section). (See also Figure S5)

Notably, we found a negative correlation between the expression of genes involved in lipid metabolism biosynthesis, modification of fatty acids, cholesterol metabolism, and phospholipid metabolism in LC2 cells with Type-II HMO concentration in milk. These genes include diacylglycerol O-acyltransferase 1 (*DGAT1*), sterol-C5-desaturase (*SC5D*), stearoyl-CoA desaturase (*SCD*), phospholipase A2 group XVI (*PLA2G16*), fatty acid desaturase 2 (*FADS2*), and 24-dehydrocholesterol reductase (*DHCR24*) (DESeq2 BH-adjusted p-value ≤ 0.05) (**Table S4**). We also saw a negative association between both Type-I and Type-II HMOs and related lactation genes such as butyrophilin subfamily 1 member A1 (*BTN1A1*), *LALBA*, casein beta (*CSN2*), and parathyroid hormone 1 receptor (*PTHR1*), which supports our conclusions from the co-expression analysis that different subsets of lactocytes may be involved in milk lipid, and HMO production (DESeq2 BH-adjusted p-value ≤ 0.05) (**Figure 3, Table S4**).^37^

Next, we asked if any of the genes that are associated with concentrations of Type-I and Type-II HMOs were enriched for biological functions in LC1 and LC2s. We performed gene set enrichment analysis (GSEA) on these lists (**Figure 6B; Supplementary Figure 5A**). The top positively enriched pathways in LC1 and LC2 cells represented gene ontology cellular processes such as *cytoplasmic translation*, *ribosomal biogenesis*, and *protein-RNA complex organization* pathways, crucial for cells engaging in large-scale protein synthesis for milk secretion (**Figure 6B**). In the LC2 cell type, we found a negative enrichment of the *vitamin transport* pathway associated with both Type-I and Type-II HMOs (**Figure 6B).** Some genes that contribute to the enrichment are ATP binding cassette subfamily G member 2 (*ABCG2*), GC vitamin D binding protein (*GC*), and folate receptor (*FOLR1*).^38^ Additionally in LC2 cells, lipid and cholesterol biosynthesis pathways were negatively enriched with increasing Type-II HMO concentrations (**Figure 6B).** Leading edge analysis highlighted the following genes: insulin-induced gene 1 (*INSIG1*), mevalonate kinase (*MVK*), 7 and 24-dehydrocholesterol reductase (*DHCR7* and *DHCR24*), delta 4-desaturase, sphingolipid 1 (*DEGS1*), prostaglandin E synthase 2 (*PTGES2*). Notably, we also observed a negative association between *response to progesterone* and Type-I HMOs in LC1s with leading genes including nuclear receptor coactivator 1 (*NCOA1*), VPS54 GARP complex subunit (*VPS54*), claudin 4 (*CLDN4*), and toll-like receptor 2 (*TLR2*) (**Figure S5A**). Taken together, this analysis could signify a specification of resources used within certain cells for either the biosynthesis of HMOs or for lipid production. Further exploration of this specification is required to clarify this relationship’s underlying mechanisms and implications for mammary gland physiology during lactation.

Next, we investigated putative transcription factors (TFs) that regulate these biological pathways to nominate candidates that may influence the biosynthesis of Type-I and Type-II HMOs in lactocytes. Overall, we found that the TFs that may regulate these processes were distinct in LC1 and LC2 cells based on this analysis (**Figure 6C**, **S5B**). In LC2s, we found a positive association of TF SMAD family member 5 (*SMAD5*) with genes positively associated with Type-II HMO concentrations (**Figure 6C).** SMAD5 regulates transforming growth factor beta (TGF-*β*) signaling in the mammary gland and is responsible for epithelial cell proliferation, differentiation, and morphogenesis in lactation.^39^ In both LC1 and LC2 cells, when considering only genes that are positively associated with type 1 HMO concentrations, we identified a positive association with ETS transcription factor ERG (*ERG*) and the transcription factor early growth response 1 (*EGR1*) (**Figure 6C, S5B).** These TFs have not been previously shown to regulate HMO biosynthesis, but have been reported to be associated with the concentrations of 2’FL, 3FL, LSTb, and LNFPII.^14^

## Discussion

The biosynthesis of HMOs in the mammary gland is difficult to map in humans given the lack of access to lactating breast samples, sufficient *ex vivo* models, and the overall complexity of HMO synthesis pathways. This study provides insights into the cellular and molecular pathways underlying HMO biosynthesis in the lactating mammary gland, highlighting the potential for lactocyte subtype specialization and gene expression regulation. We addressed this challenge by coupling scRNA-seq data from human milk-derived cells with LC-MS/MS quantitation of HMO concentrations in matched human milk samples. We identified associations between cell types, genes, and pathways that could support HMO biosynthesis during human lactation. Our work enhances the understanding of HMO biosynthesis, suggesting not only pathways that may support their production, but that specific subsets of lactocytes may play divergent and possibly complimentary roles in this process during lactation. In particular, we suggest candidate glycosyltransferase genes involved in LSTb synthesis using co-expression analysis, providing insights into the molecular pathways underlying HMO production. Our work has also assembled an integrated resource of scRNA-seq of human milk-derived cells that provides a comprehensive look into the cellular composition and dynamics of human lactocytes during lactation. We believe that the utility of this resource extends beyond HMO research to other milk components or cellular programs, serving as a valuable tool for investigating the diverse facets of mammary gland biology and lactation such as identifying the roles and populations of immune cells.

We found that lipid production gene expression was negatively correlated with Type-II HMO concentration, possibly suggesting the specialization of lactocyte subtypes in lactation. This is an attractive model from the standpoint that the uniformly expressed *LALBA* and *B4GALT1* might maintain consistent levels of lactose in all lactocytes, while more specialized lactocytes may ‘fine-tune’ the HMO and lipid contents based on the cadre of genes expressed in each cell. This also may provide insights into how these cells balance their individual metabolic demands with the diverse biosynthetic demands of milk production. Our findings further revealed many genes whose expression correlated with HMO concentrations that were specific to either LC1 or LC2 cell types, with little overlap, suggesting unique synthesis steps used by each cell type. Cell type specialization underpins the overall function of multi-cellular organisms, and our results provide evidence for this lactocyte subtype specialization in the context of the lactating mammary gland.^40^ As logistical challenges preclude the use of biopsies of lactating mammary tissue, milk fat globules, which comprise lipid droplets and cytoplasmic fragments encapsulated by plasma membrane, are commonly used as a source of lactocyte cytoplasmic material.^41^ Our findings suggest that the cells which secrete milk fat globules may not be the same cells that synthesize HMOs, and therefore, milk fat globules may not provide an accurate representation of HMO synthesizing cells.

There were several HMOs, including LSTb, DSLNT, DFLNT, LNFPI, and LNFPIII, that were positively associated with the expression of *ST6GAL1* in LC1 cells; LC2 cells, in contrast, exhibited positive associations with these HMOs and expression of *ST3GAL1*. There is no previously hypothesized role for these sialyltransferases in the production of these five HMOs except for the HMO DFLNT, where the presence of an N-acetylneuraminic acid suggests a role for a member of the *ST3GAL* family in an upstream synthesis step (**Figure 1A**). Our analysis also supports the existing hypothesis that HMO synthesis and transport occur through an ER-Golgi network. We saw positive associations between specific HMOs and genes involved in ER transport, and Golgi trafficking through the COPI pathway, specifically *COPB2* and LNFPI. The COPI pathway is involved in the secretory biosynthesis, transport, and packaging of milk components, like proteins, lipids, and carbohydrates. These findings suggest a specialized transport pathway for HMOs that warrants further investigation to understand the process of HMO biosynthesis and secretion.

While our study offers valuable insights into the association between HMO concentrations and cellular dynamics, a few limitations should be acknowledged. Firstly, glycosyltransferase genes are not highly expressed in milk-derived lactocytes. Thus, their detection may be challenging and does not directly represent protein abundances, potentially leading to the underrepresentation of certain molecular pathways. Additionally, our correlative analyses limit our ability to infer direct relationships between variables. The lack of association between hypothesized HMO synthesis gene expression and HMO concentrations could be attributed to the possibility that a smaller, more specialized group of HMO-producing cells exists within one of these LC1 or LC2 populations. The specific expression of these genes by this specialized group may be hidden by other similar cells in this aggregated pseudobulk analysis. Alternatively, a direct relationship between RNA and protein expression levels may be lacking. We attempted to cluster cells based on high expression of known HMO synthesis genes; however, the heterogeneity of these epithelial cells, their diverse functions, and the low expression of these HMO synthesis genes impaired our ability to identify cohesive cell subsets involved in these processes. Furthermore, the assumption that changes in the proportion of LC1 and LC2 cells reflect alterations in mammary gland activity may be confounded by factors such as pumping or weaning practices, warranting a cautious interpretation of studies using milk-derived cells. Finally, our findings on associations between immune cell proportions and HMO concentrations could point towards some immunomodulatory properties of HMOs or could be the result of some underlying covariate that we are unable to account for, the fact that immune cell types are rare in milk, or that the cells in the milk do not necessarily represent the composition of cells resident in the mammary gland.

Our analyses have revealed consistent differences between LC1 and LC2 cells, suggesting distinct roles in HMO production and secretion. Future work should seek to determine whether these factors are cell-intrinsic (e.g., the underlying programs of LC1 and LC2 cells) or are governed by interactions within the tissue niche or by the physiological demands of lactation. Interestingly, a greater number of genes and pathways were associated with HMO concentrations in LC2 cells, indicating a potentially more intricate regulatory network governing HMO biosynthesis in these cells. Moving forward, future research efforts that prioritize the development of functional models for human milk production will enable more comprehensive investigations into the mechanisms governing HMO biosynthesis.

## Methods

### scRNA-seq data sources

Data were downloaded from prior studies Nyquist et al (Single Cell Portal https://singlecell.broadinstitute.org/single_cell/study/SCP1671), Twigger et al (Array Express database at following access IDs: “E-MTAB-9841” (Batch 1), “E-MTAB-10855” (Batch 2) and “E-MTAB-10885” (Batch 3)) and Martin Carli et al (GEO accession GSE153889).

### Harmony for data integration

We performed data integration using pyharmony through the Scanpy function scanpy.external.pp.harmony_integrate with default parameters and batch set to ‘study’.^42,43^ Following integration, Leiden clustering was used to re-cluster the cells, identifying 14 clusters which were then manually classified into LC1, LC2, cycling epithelial, myeloid, and lymphoid cell clusters based on marker genes from the source studies. We used the same approach for integration with mammary gland tissue samples resulting in three additional clusters of resting mammary gland cells.

### DE between LC1 and LC2 across studies

Using the DESeq2 on per-sample pseudobulk counts of LC2 and LC1 cell types, we performed differential expression analysis between LC1 and LC2 cell types across samples and studies. Pseudobulk counts were aggregated by summed counts within each biological sample from cell types identified in the integrated data across studies. DESeq2 was then run using a Wald test with the model formula ‘∼ Study+cluster’ to identify genes reproducibly differential between LC1 and LC2 clusters.^25^

### Gene-gene co-expression analysis

A hypergeometric test was run on each candidate HMO gene pair in each sample in LC1 and LC2 cell types using the Python Scipy package function *scipy.stats.hypergeom* with the total number of cells in a given cell type and sample set as the population size, the possible successes set as the number of cells for which gene expression of the first gene was greater than or equal to one, the number of draws as the number of cells for which the gene expression of the second gene was expressed at a value greater than or equal to 1, and the observed successes set to the number of cells for which both genes were expressed at levels greater than or equal to one. The test was run in a given sample as long as at least 2% of cells in the given cell type expressed each gene and that the number of cells expressing each gene was at least 5. This led to a different total number of samples for which the hypergeometric test was run for each gene pair. These results were visualized showing the total number of samples for which the hypergeometric test was run and showing the percentage of these samples for which this association met the p ≤ 0.05 threshold. Clusters were identified on the top 200 gene pairs using the Seaborn clustermap function with the linkage method set to “median”.

### HPLC HMO profiling

The non-HMO oligosaccharide maltose was added to each sample as an internal standard prior to sample processing.^44^ Samples were lyophilized using a speed-vacuum before glycans were labeled with 2-aminobenzamide (2AB) at 65°C for 2 h. 2AB-glycans were separated by high-performance-liquid-chromatography with fluorescent detection (HPLC-FL) (Dionex Ultimate 3000, Dionex, now Thermo) on an TSKgel amide-80 column (15 cm length, 2 mm inner diameter, 3 μm particle size; Tosoh Bioscience). Annotation of peaks was based on standard retention times. The absolute concentration of the following HMOs was calculated based on the area under the curve for the internal standard maltose and the reference standards of each of the individual HMOs: 2’-fucosyllactose (2’FL), 3 FL, 3’SL, 6-sialyllactose (6’SL), difucosyllactose (DFLac), difucosyllacto-N-hexaose (DFLNH), difucosyllacto-N-tetrose (DFLNT), disialyllacto-N-hexaose (DSLNH), disialyllacto-N-tetraose (DSLNT), fucodisialyllacto-N-hexaose (FDSLNH), fucosyllacto-N-hexaose (FLNH), LNFP1, LNFP2, LNFP3, lacto-N-hexaose (LNH), lacto-N-tetrose (LNT), lacto-N-neo-tetrose (LNnT), sialyllacto-N-tetraose (LST) b, and LSTc. HMO-bound sialic acid (Sia) and HMO-bound fucose were calculated on a molar basis (nmol/mL).

### Mixed linear models

Linear mixed models were run with the Python package statsmodels using the function *statsmodels.formula.api.mixedlm* with fit method ‘lbfgs’.^45^ First, HMO concentrations, time postpartum, and cell-type proportions were centered and scaled by dividing by the mean and subtracting the standard deviation. To identify associations between HMO concentration and time postpartum model formula “HMO concentration ∼time postpartum” was used for each individual HMO with the groups parameter set to donor. To identify the association between HMO concentration and cell type proportion with the model formula “HMO concentration ∼time postpartum + cell type proportion” was used for each HMO and cell-type with the groups parameter set to donor. P values were corrected using Benjamini-Hochberg correction for each cell type and for the association with time using the *statsmodels.stats.multitest.multipletests* function with alpha=0.05.

### DESeq2 regression model

DESeq2 was used to identify differentially expressed genes associated with individual HMOs and Type-I and Type-II HMO concentrations. First, we created a pseudobulk of LC1 and LC2 cell populations. We had 25 samples of LC2 and 26 samples of LC1. When implementing DESeq2, we scaled and centered HMO concentrations and time postpartum variables. Based on prior analyses, we incorporated both donor identity and time postpartum as variables.^17^ We created a reduced model with only donor and time postpartum, ‘∼ donor+time_post_partum_days’, and used the likelihood ratio test to remove the effects of these covariates, isolating genes correlated with HMO concentration within LC1 and LC2 cell types. P values were corrected using Benjamini-Hochberg correction for each cell type with alpha=0.05.

### GSEA

GSEA was implemented with differentially expressed gene lists, which were the outputs from DESeq2 analysis. To rank the gene lists, we utilized a combination of the sign of fold change and the negative log10-transformed p-value derived from DESeq2 analysis. This ranking strategy facilitated the identification of genes that exhibited both substantial fold changes and statistical significance. Additionally, to account for multiple hypothesis testing, we applied the Benjamin-Hochberg correction method. Leading-edge analysis was used to identify a subset of genes that contribute most to the enrichment of a given pathway. Briefly, it is an assessment of overlap that exists between the leading edge genes of different sets, revealing which genes are overrepresented and consistently contribute to the enrichment across multiple gene sets.

### CHEA analysis

To identify potential transcription factors (TFs) regulating the differentially expressed genes (DEGs) in our study, we utilized the ChEA3 (ChIP Enrichment Analysis 3) platform, analyzing upregulated and downregulated genes separately, each with a (DeSeq-2 adjusted P-value < 0.05). The DEGs were divided into upregulated and downregulated sets, and upregulated genes were used as inputs in ChEA3. First, using the Fisher Exact Test (FET) significance of our input gene list was calculated against TF-gene sets across six different libraries in the CHEA3 database. Using the Benjamini–Hochberg correction method, these p-values were corrected for multiple comparisons and adjusted. Based on these corrected p-values, an integer rank is assigned to TFs, where 1 is the lowest p-value (most significant TF) up to rank k (highest p-value), for k TFs. For libraries containing multiple gene sets corresponding to the same TF, only the gene set with the lowest P-value was considered. The MeanRank method was implemented where a score is assigned to each TF, this score is calculated by taking the average of all ranks of each TF across the 6 different libraries. The number of differentially upregulated genes associated with Type-II HMO concentration in LC1s was extremely low. Hence TF analysis was not performed for this association.

## Data availability

scRNA-seq data used for integration can be found in the repositories of the original studies: Nyquist et al https://singlecell.broadinstitute.org/single_cell/study/SCP1671), Twigger et al (Array Express database at following access IDs: “E-MTAB-9841” (Batch 1), “E-MTAB-10855” (Batch 2) and “E-MTAB-10885” (Batch 3)) and Martin Carli et al (GEO accession GSE153889). HMO concentration data is available in **Supplementary Table 2.** Analysis scripts available at https://github.com/snyquist2/scRNA_HMO_HM.

## Contributions

Conceptualization: JMC, LB, AJT. Analysis: SKN, LDA, GDT, JMC, BAG. Funding: LB, BAG, BEE. Data collection: BAG, SKN, (Milk samples), KS, AF, LB (HMO profiling). Paper Writing first draft: LDA, SKN. Figure generation: LDA, SKN, JMC, BAG. Draft reviewing and editing: SKN, LDA, JMC, MR, AJT, LB, BAG, MCR, BEE. Supervision: LB, BAG, BEE

## Acknowledgments

We would like to thank Alex Shalek and Shalek lab members for supporting the original sample collection. We also would like to thank Archit Verma and Engelhardt group members for helpful analysis advice and discussion. B.A.G is supported in part by the Geisel School of Medicine at Dartmouth’s Center for Quantitative Biology through a grant from the National Institute of General Medical Sciences (NIGMS, P20GM130454) of the NIH. L.B. is the UC San Diego Chair of Collaborative Human Milk Research, endowed by the Family Larsson-Rosenquist Foundation in Switzerland. A.J.T. is funded by a Future Leaders Fellowship from the United Kingdom Research and Innovation (UKRI, MR/X035727/1). B.E.E is funded by NIH NHGRI R01 HG012967 and NIH NCI 5U2CCA233195, and she is a CIFAR Fellow in the Multiscale Human Program. J.F.M.C. is supported in part by a K99 award (K99 HD107496) and S.K.N is supported by an F32 (F32 HD114427).

## Declaration of generative AI and AI-assisted technologies

During the preparation of this work, the author(s) used chatGPT in order to edit early drafts for grammatical correctness and aid in debugging R code. After using this tool or service, the author(s) reviewed and edited the content as needed and take(s) full responsibility for the content of the publication.

## Supplemental Information

Document S1: Figures S1-5

Table S1. Excel file containing additional data too large to fit in a PDF, related to Figure 2. Mean gene expression of putative HMO biosynthesis genes in each study.

Table S2. Excel file containing additional data too large to fit in a PDF, related to Figures 4 and S2. HMO concentration values.

Table S3. Excel file containing additional data too large to fit in a PDF, related to Figures 4 and S2. Linear mixed model results of time and cell type proportion-associated HMO concentrations.

Table S4. Excel file containing additional data too large to fit in a PDF, related to Figure 5 and Table 1. DESeq2 results of gene expression in LC1 and LC2 cells with each HMO and Type I and Type II HMOS.

Table S5. Excel file containing additional data too large to fit in a PDF, related to Figure 6. GSEA results on genes associated with each HMO in LC1 or LC2 cells.

