## Supplementary material for "Integrated ‘omics analysis reveals human milk oligosaccharide biosynthesis programs in human lactocytes": Document S1

**Supplementary Information**

### Title: Integrated 'omics analysis reveals putative genes and gene programs involved in human milk oligosaccharide production in human lactocytes

Sarah Kate Nyquist[+*1], Laasya Devi Annepureddy[+2], Kristija Sejane[3], Annalee Furst[3], G Devon Trahan[4], Michael C Rudolph[5], Alecia Jane Twigger[6,7], Lars Bode[3], Barbara E Engelhardt[1,8], Jayne F Martin Carli*[9], Britt Anne Goods*[,2,10]

Affiliations:
[1] Gladstone Institute of Data Science & Biotechnology, San Francisco, CA, USA.
[2] Thayer School of Engineering Dartmouth College, Hanover, NH, USA.
[3] Department of Pediatrics, Larsson-Rosenquist Foundation Mother-Milk-Infant Center of Research Excellence (MOMI CORE), and the Human Milk Institute (HMI), University of California San Diego, La Jolla, CA
[4] Department of Pediatrics, Section of Hematology, Oncology, and Bone Marrow Transplant, University of Colorado Anschutz Medical Campus, Aurora, CO, USA.
[5] Harold Hamm Diabetes Center, Department of Biochemistry and Physiology, University of Oklahoma Health Sciences Center, Oklahoma City, OK, USA.
[6] Department of Pharmacology, University of Cambridge, Cambridge, England; Department of
[7] Biochemistry, University of Cambridge, Cambridge, England.
[8] Department of Biomedical Data Science, Stanford University, Stanford, USA
[9] Department of Obstetrics and Gynecology, Division of Reproductive Sciences, University of Colorado Anschutz Medical Campus, Aurora, CO, USA.
[10] Department of Molecular and Systems Biology, and Program in Quantitative Biomedical Sciences at Dartmouth College, Hanover, NH, USA.

[*]Equal contributions corresponding authors
[+]Equal contributions

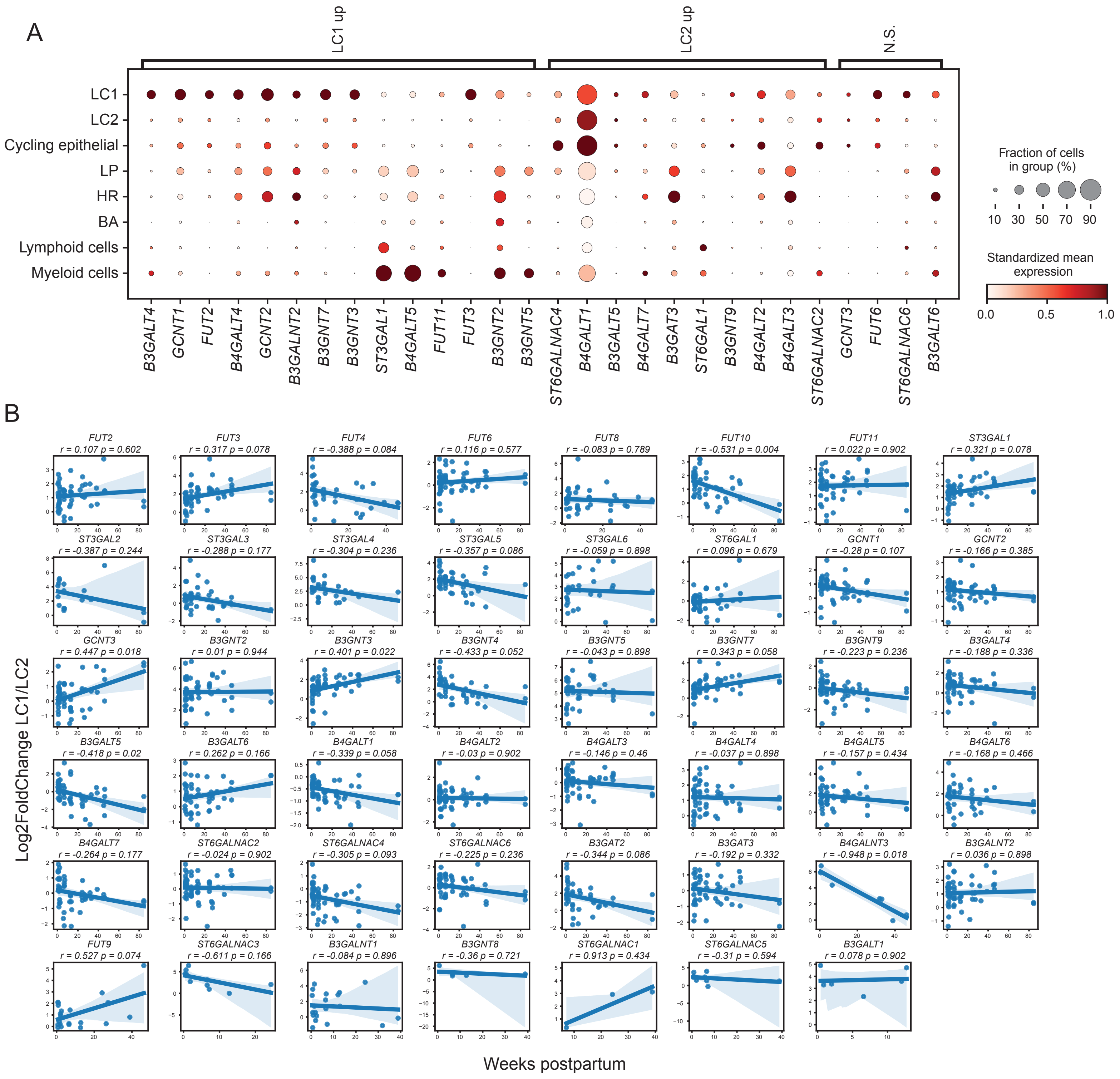

A
LC1 up
LC2 up
N.S.
LC1
LC2
Cycling epithelial
LP
HR
BA
Lymphoid cells
Myeloid cells
B3GALT4
GCNT1
FUT2
B4GALT4
GCNT2
B3GALNT2
B3GNT7
B3GNT3
ST3GAL1
B4GALT5
FUT11
FUT3
B3GNT2
B3GNT5
ST6GALNAC4
B4GALT1
B3GALT5
B4GALT7
B3GAT3
ST6GAL1
B3GNT9
B4GALT2
B4GALT3
ST6GALNAC2
GCNT3
FUT6
ST6GALNAC6
B3GALT6
Fraction of cells in group (%)
10 30 50 70 90
Standardized mean expression
0.0 0.5 1.0
B
FUT2 r = 0.107 p = 0.602
FUT3 r = 0.317 p = 0.078
FUT4 r = -0.388 p = 0.084
FUT6 r = 0.116 p = 0.577
FUT8 r = -0.083 p = 0.789
FUT10 r = -0.531 p = 0.004
FUT11 r = 0.022 p = 0.902
ST3GAL1 r = 0.321 p = 0.078
ST3GAL2 r = -0.387 p = 0.244
ST3GAL3 r = -0.288 p = 0.177
ST3GAL4 r = -0.304 p = 0.236
ST3GAL5 r = -0.357 p = 0.086
ST3GAL6 r = -0.059 p = 0.898
ST6GAL1 r = 0.096 p = 0.679
GCNT1 r = -0.28 p = 0.107
GCNT2 r = -0.166 p = 0.385
GCNT3 r = 0.447 p = 0.018
B3GNT2 r = 0.01 p = 0.944
B3GNT3 r = 0.401 p = 0.022
B3GNT4 r = -0.433 p = 0.052
B3GNT5 r = -0.043 p = 0.898
B3GNT7 r = 0.343 p = 0.058
B3GNT9 r = -0.223 p = 0.236
B3GALT4 r = -0.188 p = 0.336
B3GALT5 r = -0.418 p = 0.02
B3GALT6 r = 0.262 p = 0.166
B4GALT1 r = -0.339 p = 0.058
B4GALT2 r = -0.03 p = 0.902
B4GALT3 r = -0.146 p = 0.46
B4GALT4 r = -0.037 p = 0.898
B4GALT5 r = -0.157 p = 0.434
B4GALT6 r = -0.168 p = 0.466
B4GALT7 r = -0.264 p = 0.177
ST6GALNAC2 r = -0.024 p = 0.902
ST6GALNAC4 r = -0.305 p = 0.093
ST6GALNAC6 r = -0.225 p = 0.236
B3GAT2 r = -0.344 p = 0.086
B3GAT3 r = -0.192 p = 0.332
B4GALNT3 r = -0.948 p = 0.018
B3GALNT2 r = 0.036 p = 0.898
FUT9 r = 0.527 p = 0.074
ST6GALNAC3 r = -0.611 p = 0.166
B3GALNT1 r = -0.084 p = 0.896
B3GNT8 r = -0.36 p = 0.721
ST6GALNAC1 r = 0.913 p = 0.434
ST6GALNAC5 r = -0.31 p = 0.594
B3GALT1 r = 0.078 p = 0.902
Log2FoldChange LC1/LC2
Weeks postpartum

Supplementary Figure 1: A. Dotplot showing expression of select candidate HMO synthesis genes (bottom) expressed in a minimum of 10 percent of cells in human milk scRNA-seq data and mammary gland tissue data grouped by celltype (left). LC1 and LC2 enrichment of each gene identified using DESeq2 comparisons of pseudo bulk data between LC1 and LC2 cells in human milk. N.S. genes have large effect size but do not reach significance in the comparison. Dot size indicates percent of cells in group expressing each gene, and dot color indicating column-standardized mean expression of each gene across cell type groups. Mammary gland tissue cell types: LP- luminal progenitor, HR- hormone responsive, BA - basal cell. B. Spearman correlation of the log2FoldChange between expression in LC1 cells and LC2 cells of potential HMO synthesis genes with time postpartum.

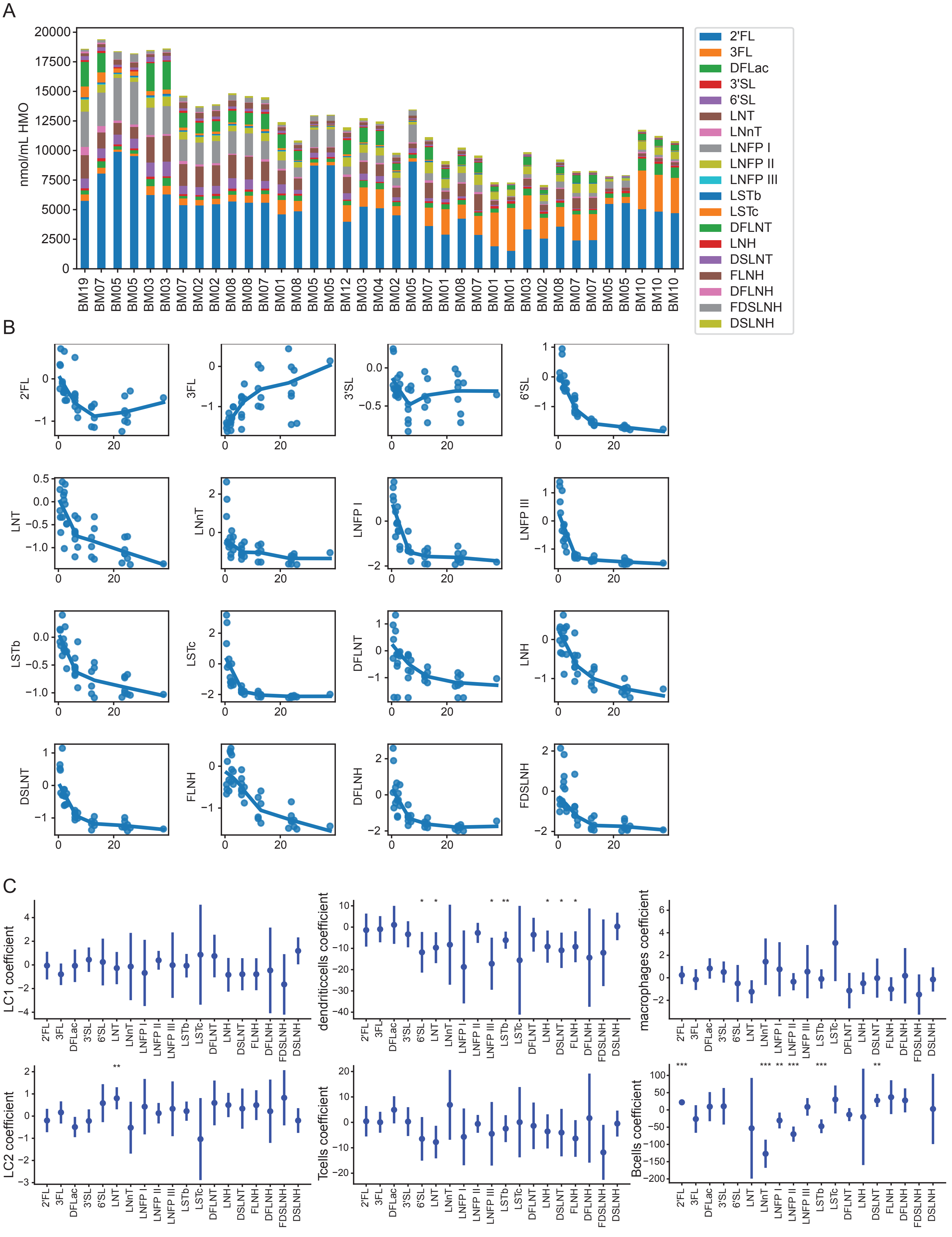

A
nmol/mL HMO
20000
17500
15000
12500
10000
7500
5000
2500
0
BM19
BM07
BM05
BM05
BM03
BM03
BM07
BM02
BM02
BM08
BM08
BM07
BM01
BM08
BM05
BM05
BM12
BM03
BM04
BM02
BM05
BM07
BM01
BM08
BM07
BM01
BM01
BM03
BM02
BM08
BM07
BM07
BM05
BM05
BM10
BM10
BM10
2'FL
3FL
DFLac
3'SL
6'SL
LNT
LNnT
LNFP I
LNFP II
LNFP III
LSTb
LSTc
DFLNT
LNH
DSLNT
FLNH
DFLNH
FDSLNH
DSLNH
B
2'FL
3FL
3'SL
6'SL
LNT
LNnT
LNFP I
LNFP III
LSTb
LSTc
DFLNT
LNH
DSLNT
FLNH
DFLNH
FDSLNH
0
20
C
LC1 coefficient
dendriticcells coefficient
macrophages coefficient
LC2 coefficient
Tcells coefficient
Bcells coefficient
2'FL
3FL
DFLac
3'SL
6'SL
LNT
LNnT
LNFP I
LNFP II
LNFP III
LSTb
LSTc
DFLNT
LNH
DSLNT
FLNH
DFLNH
FDSLNH
DSLNH
*
**
***

Supplementary Figure 2: A. Sample composition of HMO concentrations ordered by increasing time postpartum, including samples with matched scRNA-seq data. For samples with multiple replicates, mean HMO concentrations are visualized.
B. As in Fig. 4B, Loess fits for significant association between HMO concentration and weeks postpartum. Scatter plots indicate HMO concentrations by sample. Significance determined by linear mixed model fits p <= 0.05.
C. Linear mixed model coefficients for association between HMO concentration and matched immune cell type proportion *p<=0.05, **p<=0.01,***p<=0.001.

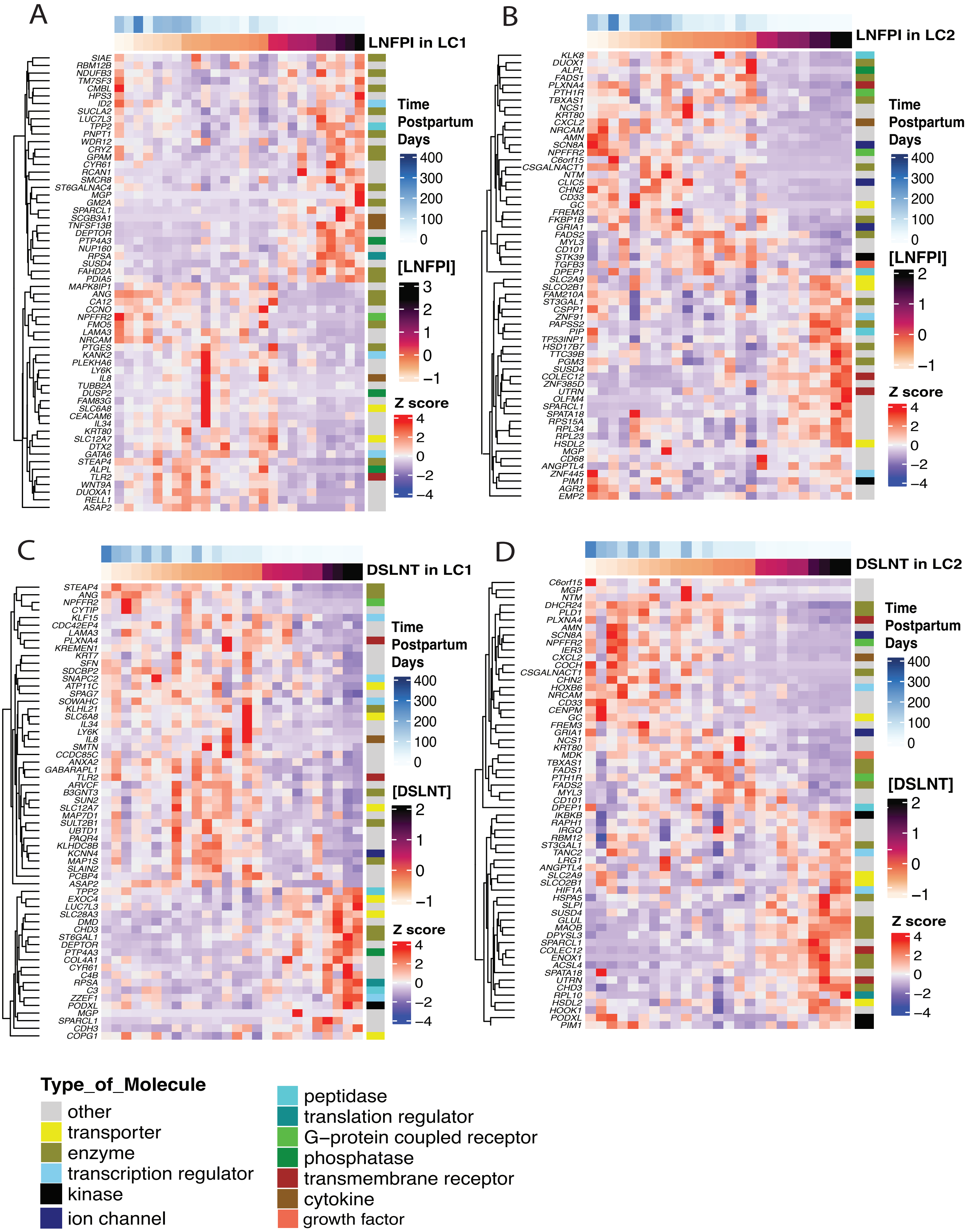
A
LNFPI in LC1
SIAE
RBM12B
NDUFB3
TM7SF3
CMBL
HPS3
ID2
SUCLA2
LUC7L3
TPP2
PNPT1
WDR12
CRYZ
GPAM
CYR61
RCAN1
SMCR8
ST6GALNAC4
MGP
GM2A
SPARCL1
SCGB3A1
TNFSF13B
DEPTOR
PTP4A3
NUP160
RPSA
SUSD4
FAHD2A
PDIA5
MAPK8IP1
ANG
CA12
CCNO
NPFFR2
FMO5
LAMA3
NRCAM
PTGES
KANK2
PLEKHA6
LY6K
IL8
TUBB2A
DUSP2
FAM83G
SLC6A8
CEACAM6
IL34
KRT80
SLC12A7
DTX2
GATA6
STEAP4
ALPL
TLR2
WNT9A
DUOXA1
RELL1
ASAP2
Time Postpartum Days
400
300
200
100
0
[LNFPI]
3
2
1
0
−1
Z score
4
2
0
−2
−4
B
LNFPI in LC2
KLK8
DUOX1
ALPL
FADS1
PLXNA4
PTH1R
TBXAS1
NCS1
KRT80
CXCL2
NRCAM
AMN
SCN8A
NPFFR2
C6orf15
CSGALNACT1
NTM
CLIC5
CHN2
CD33
GC
FREM3
FKBP1B
GRIA1
FADS2
MYL3
CD101
STK39
TGFB3
DPEP1
SLC2A9
SLCO2B1
FAM210A
ST3GAL1
CSPP1
ZNF91
PAPSS2
PIP
TP53INP1
HSD17B7
TTC39B
PGM3
SUSD4
COLEC12
ZNF385D
UTRN
OLFM4
SPARCL1
SPATA18
RPS15A
RPL34
RPL23
HSDL2
MGP
CD68
ANGPTL4
ZNF445
PIM1
AGR2
EMP2
Time Postpartum Days
400
300
200
100
0
[LNFPI]
2
1
0
−1
Z score
4
2
0
−2
−4
C
DSLNT in LC1
STEAP4
ANG
NPFFR2
CYTIP
KLF15
CDC42EP4
LAMA3
PLXNA4
KREMEN1
KRT7
SFN
SDCBP2
SNAPC2
ATP11C
SPAG7
SOWAHC
KLHL21
SLC6A8
IL34
LY6K
IL8
SMTN
CCDC85C
ANXA2
GABARAPL1
TLR2
ARVCF
B3GNT3
SUN2
SLC12A7
MAP7D1
SULT2B1
UBTD1
PAQR4
KLHDC8B
KCNN4
MAP1S
SLAIN2
PCBP4
ASAP2
TPP2
EXOC4
LUC7L3
SLC28A3
DMD
CHD3
ST6GAL1
DEPTOR
PTP4A3
COL4A1
CYR61
C4B
RPSA
C3
ZZEF1
PODXL
MGP
SPARCL1
CDH3
COPG1
Time Postpartum Days
400
300
200
100
0
[DSLNT]
2
1
0
−1
Z score
4
2
0
−2
−4
D
DSLNT in LC2
C6orf15
MGP
NTM
DHCR24
PLD1
PLXNA4
AMN
SCN8A
NPFFR2
IER3
CXCL2
COCH
CSGALNACT1
CHN2
HOXB6
NRCAM
CD33
CENPM
GC
FREM3
GRIA1
NCS1
KRT80
MDK
TBXAS1
FADS1
PTH1R
FADS2
MYL3
CD101
DPEP1
IKBKB
RAPH1
IRGQ
RBM12
ST3GAL1
TANC2
LRG1
ANGPTL4
SLC2A9
SLCO2B1
HIF1A
HSPA5
SLPI
SUSD4
GLUL
MAOB
DPYSL3
SPARCL1
COLEC12
ENOX1
ACSL4
SPATA18
UTRN
CHD3
RPL10
HSDL2
HOOK1
PODXL
PIM1
Time Postpartum Days
400
300
200
100
0
[DSLNT]
2
1
0
−1
Z score
4
2
0
−2
−4
Type_of_Molecule
other
transporter
enzyme
transcription regulator
kinase
ion channel
peptidase
translation regulator
G−protein coupled receptor
phosphatase
transmembrane receptor
cytokine
growth factor

**Supplementary Figure 3**: A. Heatmap of genes associated with the concentration of LNFPI in LC1 cell type. B. Heatmap of genes associated with the concentration of LNFPI in LC2 cell type. C. Heatmap of genes associated with the concentration of DSLNT in LC1 cell type. D. Heatmap of genes associated with the concentration of DSLNT in LC2 cell type. Heatmap includes z-scored mean expression of pseudo bulk counts within each sample's LC1 (left) or LC2 (right) ordered by HMO concentration and annotated by time postpartum and type of molecule from the IPA database. Genes are hierarchically clustered using Euclidean distance.

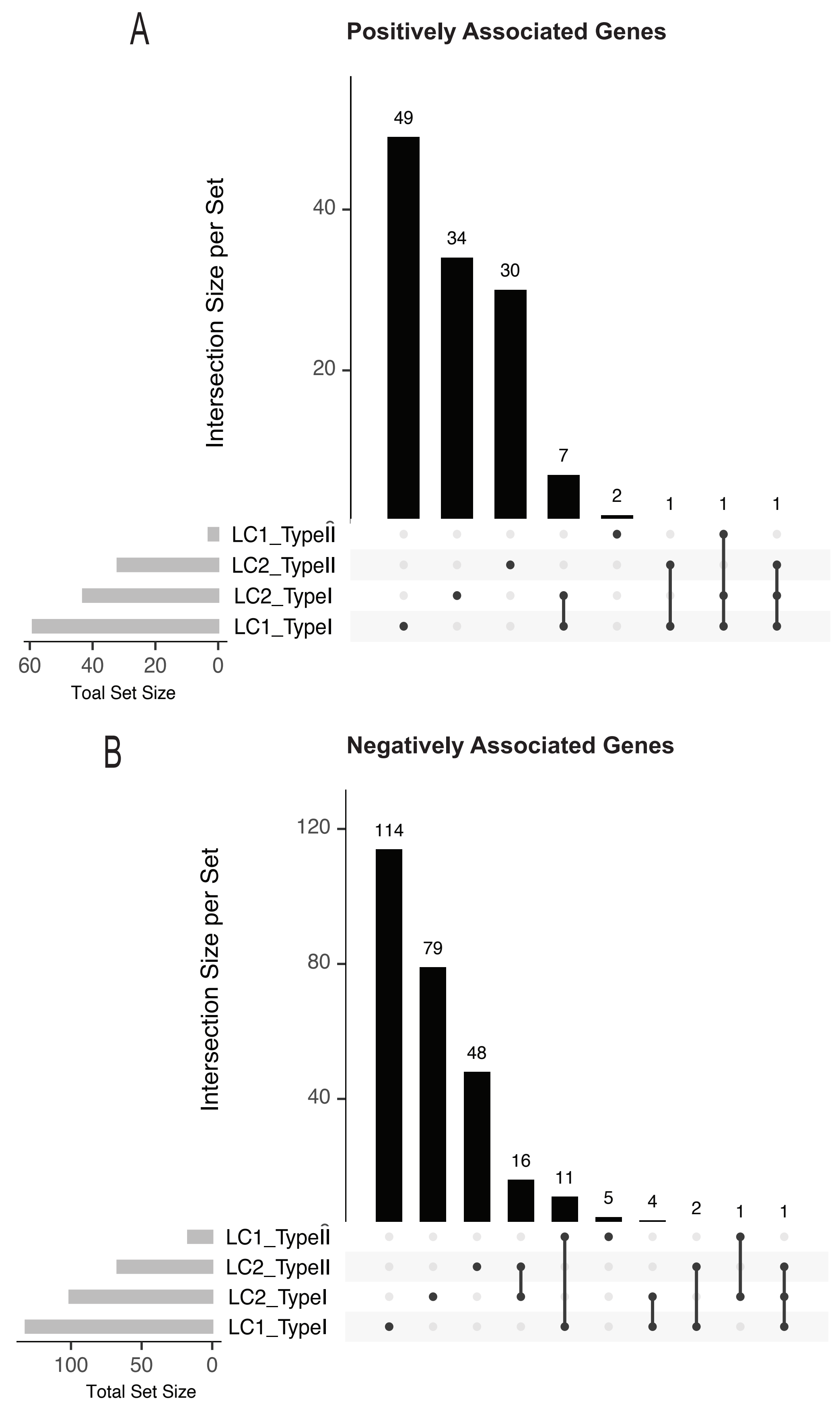
A
Positively Associated Genes
Intersection Size per Set
40
20
49
34
30
7
2
1
1
1
LC1_TypeII
LC2_TypeII
LC2_TypeI
LC1_TypeI
60
40
20
0
Toal Set Size
B
Negatively Associated Genes
Intersection Size per Set
120
80
40
114
79
48
16
11
5
4
2
1
1
LC1_TypeII
LC2_TypeII
LC2_TypeI
LC1_TypeI
100
50
0
Total Set Size

**Supplementary Figure 4:** UpSet plot showing the number of uniquely associated or intersecting genes (vertical bars) which were associated with Type I and Type II HMO concentration in LC1 and LC2 cell type. The horizontal bars indicate total number of genes in the set. A. UpSet plot only for upregulated genes. B. UpSet plot only for downregulated genes (bottom). DESeq2 BH adjusted p-value = <0.05 for all genes.

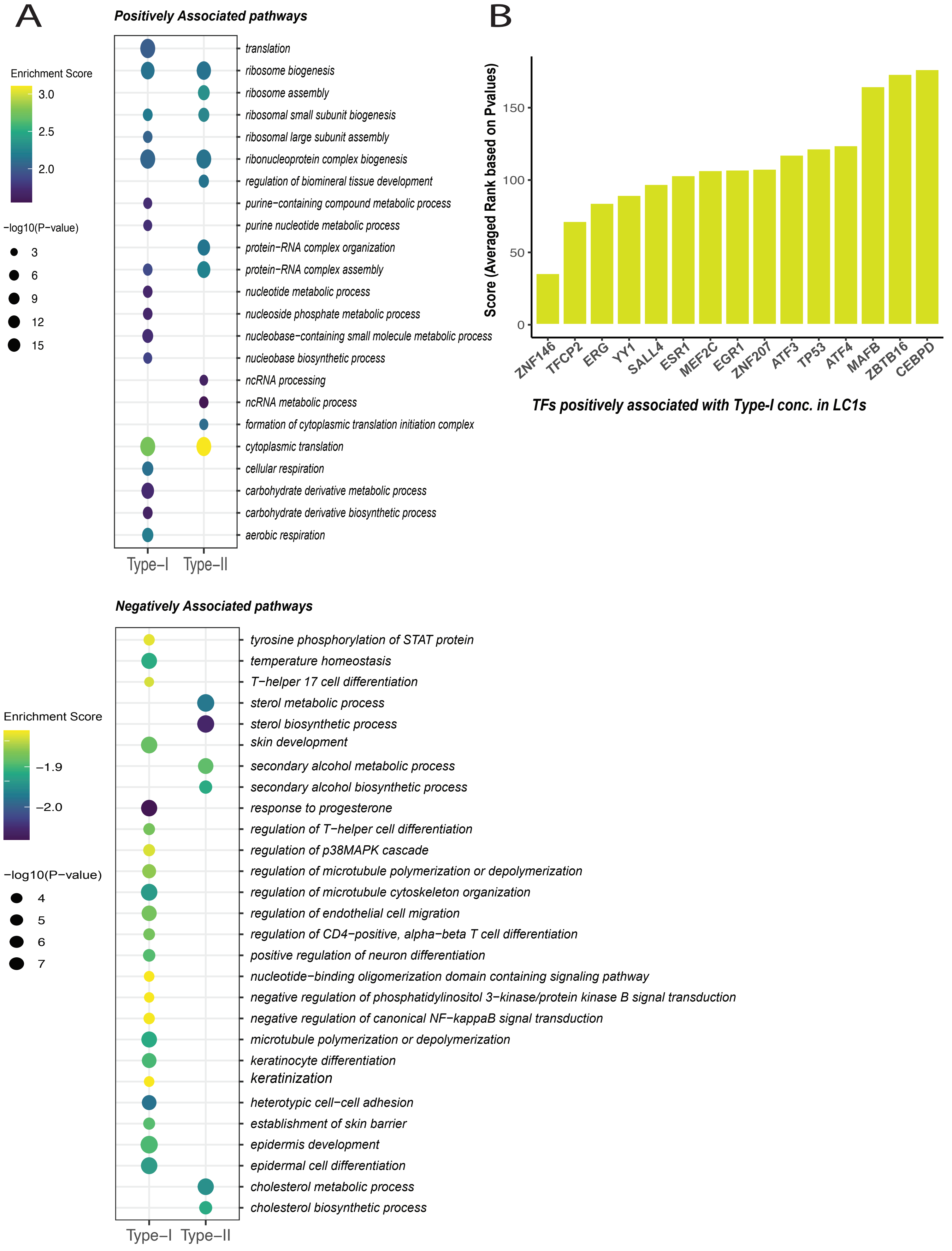
A
Positively Associated pathways
Enrichment Score
3.0
2.5
2.0
−log10(P−value)
3
6
9
12
15
translation
ribosome biogenesis
ribosome assembly
ribosomal small subunit biogenesis
ribosomal large subunit assembly
ribonucleoprotein complex biogenesis
regulation of biomineral tissue development
purine−containing compound metabolic process
purine nucleotide metabolic process
protein−RNA complex organization
protein−RNA complex assembly
nucleotide metabolic process
nucleoside phosphate metabolic process
nucleobase−containing small molecule metabolic process
nucleobase biosynthetic process
ncRNA processing
ncRNA metabolic process
formation of cytoplasmic translation initiation complex
cytoplasmic translation
cellular respiration
carbohydrate derivative metabolic process
carbohydrate derivative biosynthetic process
aerobic respiration
Type−I
Type−II
B
Score (Averaged Rank based on Pvalues)
0
50
100
150
ZNF146
TFCP2
ERG
YY1
SALL4
ESR1
MEF2C
EGR1
ZNF207
ATF3
TP53
ATF4
MAFB
ZBTB16
CEBPD
TFs positively associated with Type-I conc. in LC1s
Negatively Associated pathways
Enrichment Score
−1.9
−2.0
−log10(P−value)
4
5
6
7
tyrosine phosphorylation of STAT protein
temperature homeostasis
T−helper 17 cell differentiation
sterol metabolic process
sterol biosynthetic process
skin development
secondary alcohol metabolic process
secondary alcohol biosynthetic process
response to progesterone
regulation of T−helper cell differentiation
regulation of p38MAPK cascade
regulation of microtubule polymerization or depolymerization
regulation of microtubule cytoskeleton organization
regulation of endothelial cell migration
regulation of CD4−positive, alpha−beta T cell differentiation
positive regulation of neuron differentiation
nucleotide−binding oligomerization domain containing signaling pathway
negative regulation of phosphatidylinositol 3−kinase/protein kinase B signal transduction
negative regulation of canonical NF−kappaB signal transduction
microtubule polymerization or depolymerization
keratinocyte differentiation
keratinization
heterotypic cell−cell adhesion
establishment of skin barrier
epidermis development
epidermal cell differentiation
cholesterol metabolic process
cholesterol biosynthetic process
Type−I
Type−II

**Supplementary Figure 5**: **A.** GSEA GO BP dotplot of top positively enriched pathways (top) and top negatively enriched pathways (bottom) associated with Type I or Type II HMO concentrations in LC1. FDR < 0.05 **B.** Top 15 transcription factors associated with upregulated DEGs associated with Type-I HMO concentration in LC1 from CHEA analysis. The mean scores are averaged ranks of each TF across multiple gene sets in the CHEA database. The mean scores are averaged ranks of each TF across multiple gene sets in the CHEA database. The rank of each TF in a gene set is assigned based on BH adjusted p-values. Lower p-values (higher significance) indicate lower integer ranks and lower overall scores (see methods section).
